# SpatialOmicsLab: an integrated research environment for AI co-scientists in spatial transcriptomics

**DOI:** 10.64898/2026.10.01.756049

**Authors:** Yi Jiang, Xiaowei Zhan, Peiran Quan, Ruheng Wang, Fangjiang Wu, Junhui Mi, Jia Yao, Bo Yao, Guanghua Xiao, Wenqi Shi, Yang Xie

## Abstract

Spatial profiling is moving from method development into a central platform for biological discovery and clinical translation, but analyzing such complex data has become the bottleneck of scientific discovery. Translating spatial transcriptomic measurements into biological insight requires reproducible workflows that integrate diverse analytical methods^1^. Although language-model agents can plan and execute multistep analyses^2^, fragmented data and software ecosystems hinder their use as reliable research assistants^3^. In this study, we introduce SpatialOmicsLab, an integrated research environment for spatial transcriptomics comprising 113 curated computational tools, 63 reference datasets and 4,614 task specifications. We also introduce ST-Coscientist, an AI co-scientist that uses this environment to assemble, execute and refine spatial transcriptomic analyses. Within SpatialOmicsLab, ST-Coscientist outperformed expert-led workflows and other scientific agents across six benchmark settings spanning spatial clustering, cell-type deconvolution and spatially variable gene detection. It constructed alignment-aware workflows for three-dimensional MERFISH data and addressed biological questions by selecting and analysing relevant reference datasets. Procedural memory increased the success rate of integrating and validating additional analytical tools from 48% to 95%. Together, SpatialOmicsLab and ST-Coscientist establish an extensible ecosystem for reproducible spatial transcriptomic research, enabling AI co-scientists to address evolving biological questions with an expanding analytical repertoire.

## Introduction

Spatial transcriptomics measures gene expression while preserving tissue organization, linking molecular states to cell identity, morphology and anatomical position^4^. The ability to resolve spatial domains, tissue boundaries, cellular neighborhoods and disease microenvironments in situ has made the technology central to studies of development, tumor ecology and translational medicine. Its rapid adoption has also driven a proliferation of platforms, producing an unusually heterogeneous data landscape. Sequencing-based platforms such as Visium, Slide-seq^5,6^, Visium HD^7^ and Stereo-seq^8^ capture the whole transcriptome at resolutions ranging from multicellular spots to subcellular bins, whereas imaging-based platforms such as MERFISH^9^, seqFISH^10,11^, Xenium^12^ and CosMx^13^ measure targeted gene panels at single-cell resolution. These platforms differ in spatial resolution, gene coverage, coordinate conventions, the availability of matched histology images and the file formats and data structures in which results are delivered. As studies expand from individual tissue sections to three-dimensional volumes and patient cohorts, extracting biological insight requires workflows that accommodate diverse measurements and integrate complementary analyses^14,15^.

The analytical ecosystem is equally fragmented. Foundational frameworks including Scanpy^16^, Squidpy^17^, Giotto^18^, SpatialData^1^, Seurat^19^ and scverse^20^ provide data structures and core operations. A growing collection of specialized methods addresses preprocessing, multi-slice alignment, spatial-domain detection, spatially variable gene (SVG) discovery, cell-type deconvolution, cell–cell communication and niche analysis^21^. These methods differ in software dependencies, computational requirements and output conventions, and may require raw or normalized counts, matched images or single-cell references^21,22^. Benchmarking further shows that method performance varies across platforms and tissues^23^, so the appropriate method depends on the data provided as well as the biological question. Combining methods requires reconciling data formats, parameters and outputs^1,20,24–29^, while continued methodological development adds to this complexity^21^. The resulting challenge is to translate this expanding methodological repertoire into coherent analyses that researchers can execute, validate and reproduce on their own data.

Large language model (LLM) agents offer a way to connect research questions with computational analysis through iterative reasoning and tool use^30–33^. Recent systems have generated and ranked research hypotheses^34^, coordinated multi-agent research teams^35^, guided experimental design and execution^36–38^, and conducted biomedical analyses using curated tool libraries^2,39,40^. In spatial biology, SpatialAgent^41^, STAgent^42^, STAT^43^, EnsAgent^44^, STAnalyzer^45^, PhenoGraph^46^ and ChatSpatial^47^ demonstrate capabilities in workflow planning, data exploration and interpretation. ChatSpatial, for example, coordinates analytical methods through schema-constrained interfaces to support reproducible analyses across platforms^47^. Despite this progress, benchmarks reveal persistent difficulties in completing realistic analytical tasks^48–57^. Performance depends on the computational environment as well as the underlying model^55^. As Luebbert observes, navigating biological data infrastructure with AI agents is “like driving through an old city that was designed before cars”^3^.

In spatial transcriptomics, method selection, execution and biological interpretation are closely coupled. An agent must choose methods that address the biological question while accounting for the measurement platform, spatial resolution, available modalities, computational resources and benchmark evidence^23^. Execution, in turn, rarely involves a single tool: most analyses chain several methods whose interfaces must be made compatible^1^. Linking spatial-domain detection with cell-type deconvolution, for example, requires reconciling inputs, parameters and outputs; selecting methods for either task also requires a common basis for comparison^1,20,24–29^. Successful execution alone is insufficient: misinterpreted coordinates, malformed outputs or silently omitted steps can compromise downstream biological inference even when code completes without error^50,55^. Agents therefore need access to input checks, intermediate results and execution feedback to scrutinize their analyses and revise subsequent decisions. Such an environment must remain extensible, so that new methods can be incorporated and tested as research needs evolve^21^, and agents can retrieve relevant reference data when none are supplied. Shared tasks and quantitative criteria are needed to assess complete workflows, from raw data to results that can be independently evaluated. Agent–computer interfaces in software engineering offer a precedent for making computational actions and their consequences accessible to agents^58^. Spatial transcriptomics calls for a corresponding research environment that provides the data, methodological context and execution feedback agents need to plan, evaluate and refine reproducible analyses^59^.

In this study, we introduce SpatialOmicsLab, an integrated research environment for spatial transcriptomics, and ST-Coscientist, an AI co-scientist that operates within it (Fig. 1). SpatialOmicsLab combines 113 computational tools (Supplementary Tables 1 and 2) and 63 curated reference datasets with standardized interfaces and isolated software environments to support reproducible analysis. Its evaluation framework comprises 4,614 task specifications (Supplementary Tables 3 and 4), together with 103 reference-annotated benchmark tasks for quantitative comparison with expert workflows and other agents (Supplementary Tables 5 and 6). Within this environment, ST-Coscientist translates biological questions into analytical workflows, selecting and configuring methods according to the data and benchmark evidence, and evaluating intermediate outputs to guide subsequent analyses. For questions posed without data, it retrieves relevant reference datasets. ST-Coscientist can also extend the environment by incorporating additional analytical methods, drawing on procedural memory to guide their integration and validation.

**Figure 1.**
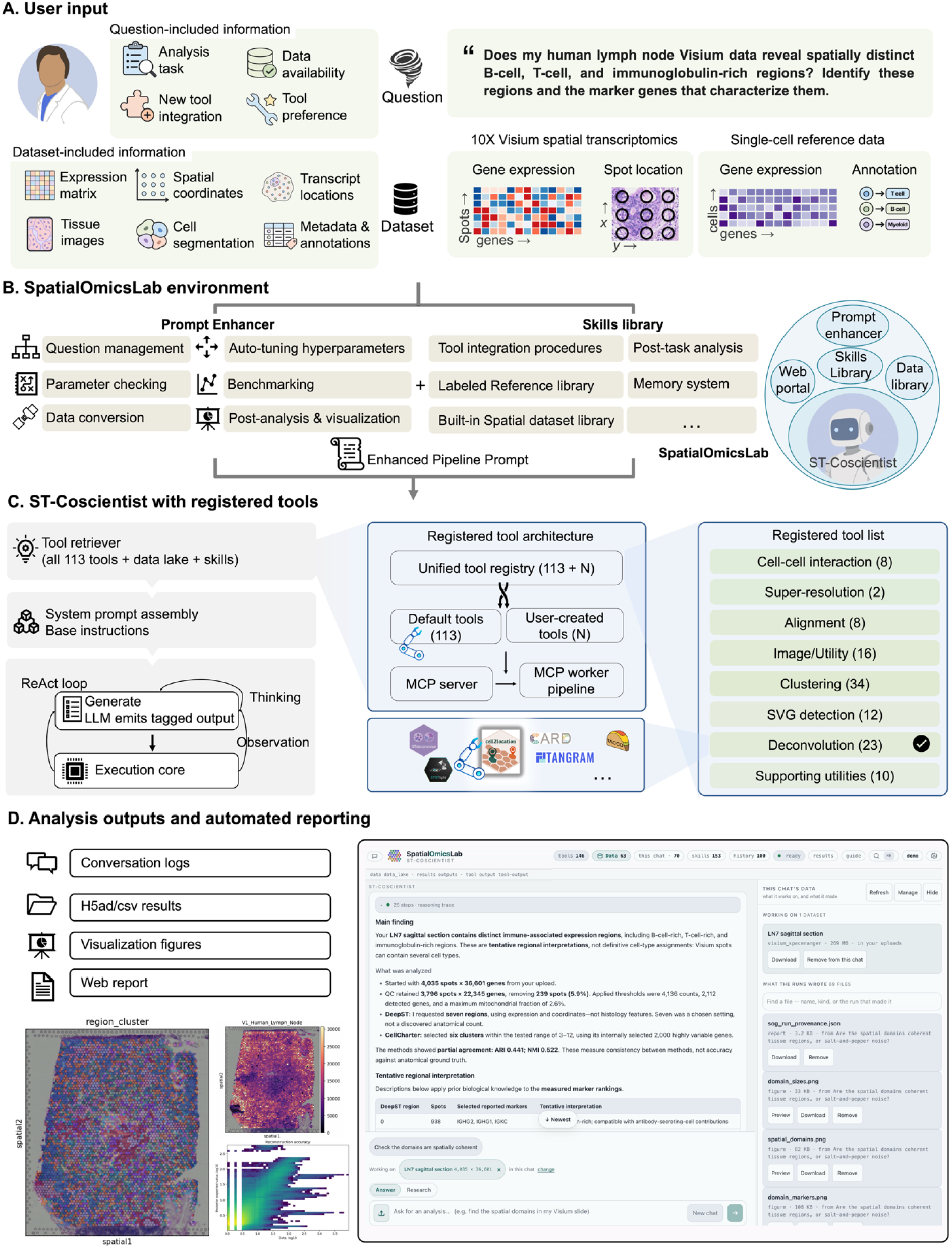
SpatialOmicsLab provides an integrated environment for spatial transcriptomic research. A, User requests and available data establish the analytical context, illustrated by a question about spatially distinct B-cell, T-cell and immunoglobulin-rich regions in human lymph-node Visium data, accompanied by a single-cell reference. B, Prompt enhancement combines data context with guidance on data conversion, parameter checking, hyperparameter tuning, benchmarking and visualization. A complementary skills library provides reference resources and reusable procedures for method integration and analysis, together with procedural memory. C, ST-Coscientist retrieves candidate tools, assembles task-specific instructions and iterates between reasoning, tool execution and observation. The registry combines 113 default tools with additional tools (*N*), accessed through Model Context Protocol (MCP) server–worker pairs. The highlighted deconvolution category and method logos illustrate tools relevant to the request in A. D, Conversation logs, standardized H5AD or CSV files, figures and web reports make analysis outputs available for inspection and reuse. The report screenshot presents a separate spatial-domain analysis. LLM, large language model; SVG, spatially variable gene.

We show that SpatialOmicsLab supports an AI co-scientist in adapting its analyses to biological questions while expanding the methods available to address them. Within this environment, ST-Coscientist outperformed expert-led workflows across six benchmark settings and achieved the highest observed success rates on the evaluated public subsets of SpatialBench^53^ and SpatialBench-Long^60^. The environment supported the construction of alignment-aware workflows for three-dimensional MERFISH data and the selection and analysis of reference datasets for questions posed without data. ST-Coscientist also extended the environment itself: procedural memory increased successful integration and validation of additional tools from 48% to 95%, without retraining the language model. Together, these results connect reproducible analysis with the accumulation of reusable methods and procedural knowledge, providing a foundation for AI co-scientists whose capabilities can develop alongside spatial transcriptomic research.

## Results

### SpatialOmicsLab couples adaptive analysis with reproducible execution

SpatialOmicsLab integrates analytical tools, reference data and execution support for spatial transcriptomic research by AI agents (Fig. 1). Its current registry contains 113 computational tools: 103 analytical tools spanning spatial clustering (34), cell-type deconvolution (23), image and utility operations (16), SVG detection (12), multi-slice alignment (8), cell–cell interaction (8) and super-resolution (2), together with 10 utilities for data conversion, dataset search, visualization and tool preparation (Supplementary Tables 1 and 2). A curated collection of 63 reference datasets supports analyses initiated from users without accompanying data. For new datasets provided by users, the environment provides structured descriptions of the measurement platform, spatial coordinates and available modalities. Standardized interfaces define each tool’s inputs, parameters and expected outputs, while Model Context Protocol (MCP) server–worker pairs provide access to isolated Python or R environments^61^. Input and output checks return structured feedback on unmet data requirements, execution errors and missing or malformed results. These descriptions, interfaces and checks provide the context and feedback agents need to select methods, assemble workflows and revise analyses. Results are returned as standardized files, figures and reports, with execution logs preserving a record of the analytical steps.

Within SpatialOmicsLab, agents pursue biological questions through analyses that are progressively refined in light of the data and intermediate results. An investigation can draw on user-supplied data or begin by retrieving relevant reference datasets from the environment (Fig. 1A). SpatialOmicsLab provides the context needed to work with these data, recording the measurement platform, file format, spatial coordinates and availability of histology images, single-cell references and metadata, with conversion to H5AD where necessary. This context is paired with task-specific guidance on parameter checking, hyperparameter tuning, benchmarking and visualization, helping agents relate the biological question and data characteristics to the available analytical methods. Reusable procedures support specialized steps such as preparing single-cell references for deconvolution and incorporating additional methods, with procedural memory making experience from earlier tool integrations available to subsequent work (Fig. 1B). Agents retrieve ranked candidate tools with supporting rationales and example calls, then assemble, execute and revise workflows (Fig. 1C). Before each step, they use the environment’s checking routines to assess input requirements and output locations; afterwards, they inspect logs and results against task- and tool-specific criteria. The resulting feedback supports refinement of both new and existing analyses, from correcting data preparation and adjusting parameters to reconsidering the choice of method.

Completed analyses return standardized results alongside records of how they were produced (Fig. 1D). H5AD or CSV files and figures are accompanied by conversation and execution logs, together with a report documenting the research request, methods, parameters, diagnostic messages and interpretation notes. These records provide a basis for scrutinizing analytical choices, reproducing computational steps and reusing results in subsequent investigations. To assess how this design supports research, we evaluated analytical accuracy and task completion across internal and external benchmarks (Figs. 2 and 3). Case studies examined how workflows adapt to user-provided data and how biological questions can be investigated using reference datasets (Figs. 4 and 5). We also assessed the integration and validation of additional methods with and without procedural memory (Fig. 6).

**Figure 2.**
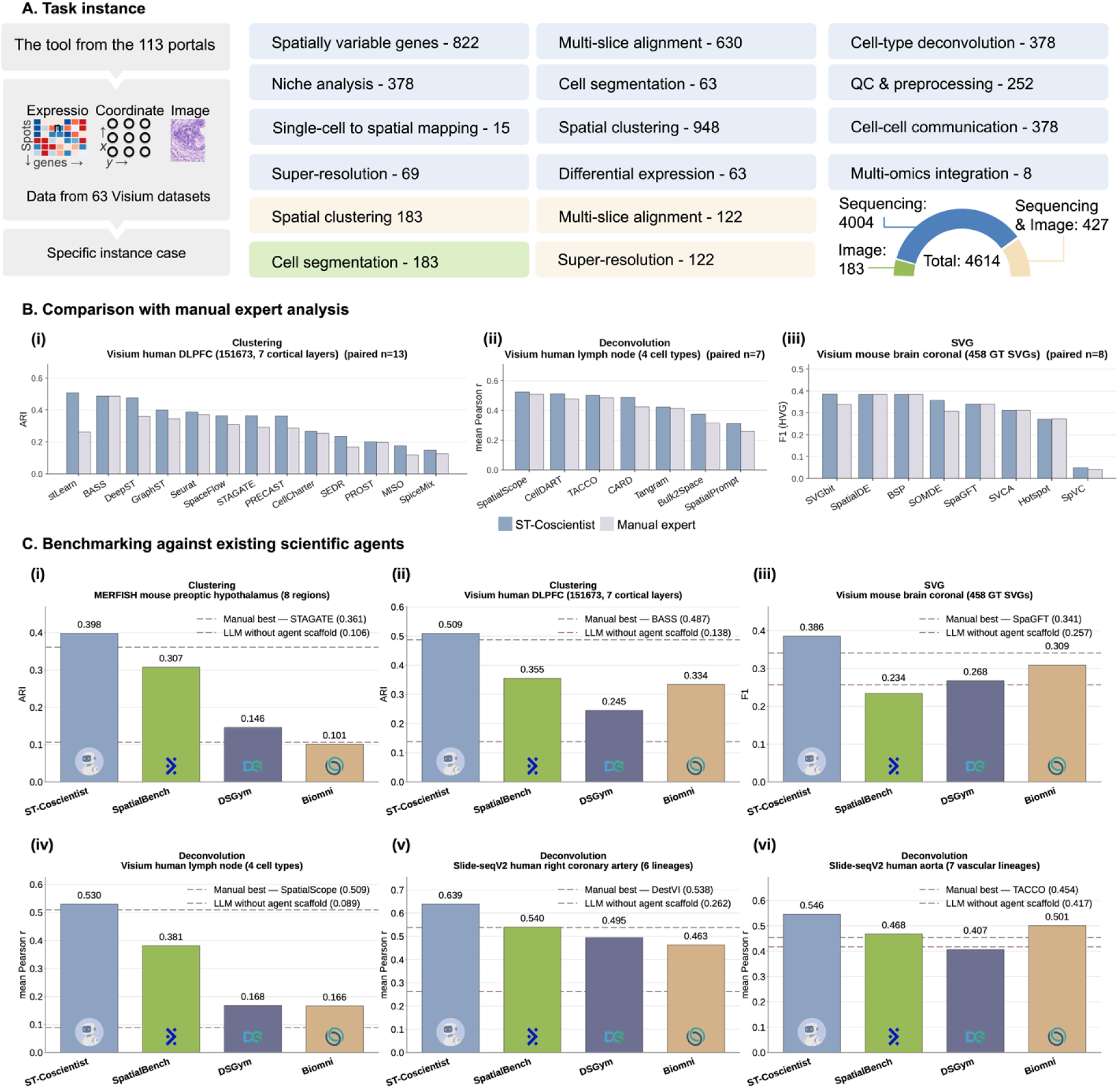
Task coverage and benchmark performance of ST-Coscientist. A, The task space contains 4,614 task specifications across 12 task families and three modalities: 4,004 sequencing-based, 183 image-based and 427 combined sequencing-and-image specifications. B, Comparisons between ST-Coscientist and expert-led workflows applying the same methods to the same data: (i) 13 spatial-clustering methods on Visium human DLPFC (sample 151673; 7 annotated tissue regions; ARI); (ii) 7 cell-type deconvolution methods on human lymph-node Visium data (4 cell types; mean Pearson r); and (iii) 8 SVG-detection methods on a Visium mouse brain coronal section (458 reference SVGs; F1). Here, n denotes the number of methods evaluated. C, Scores for ST-Coscientist and the configurations labelled SpatialBench, DSGym and Biomni across six benchmark settings. Dashed lines indicate the corresponding expert reference score and the backbone language model without an agent scaffold.

**Figure 3.**
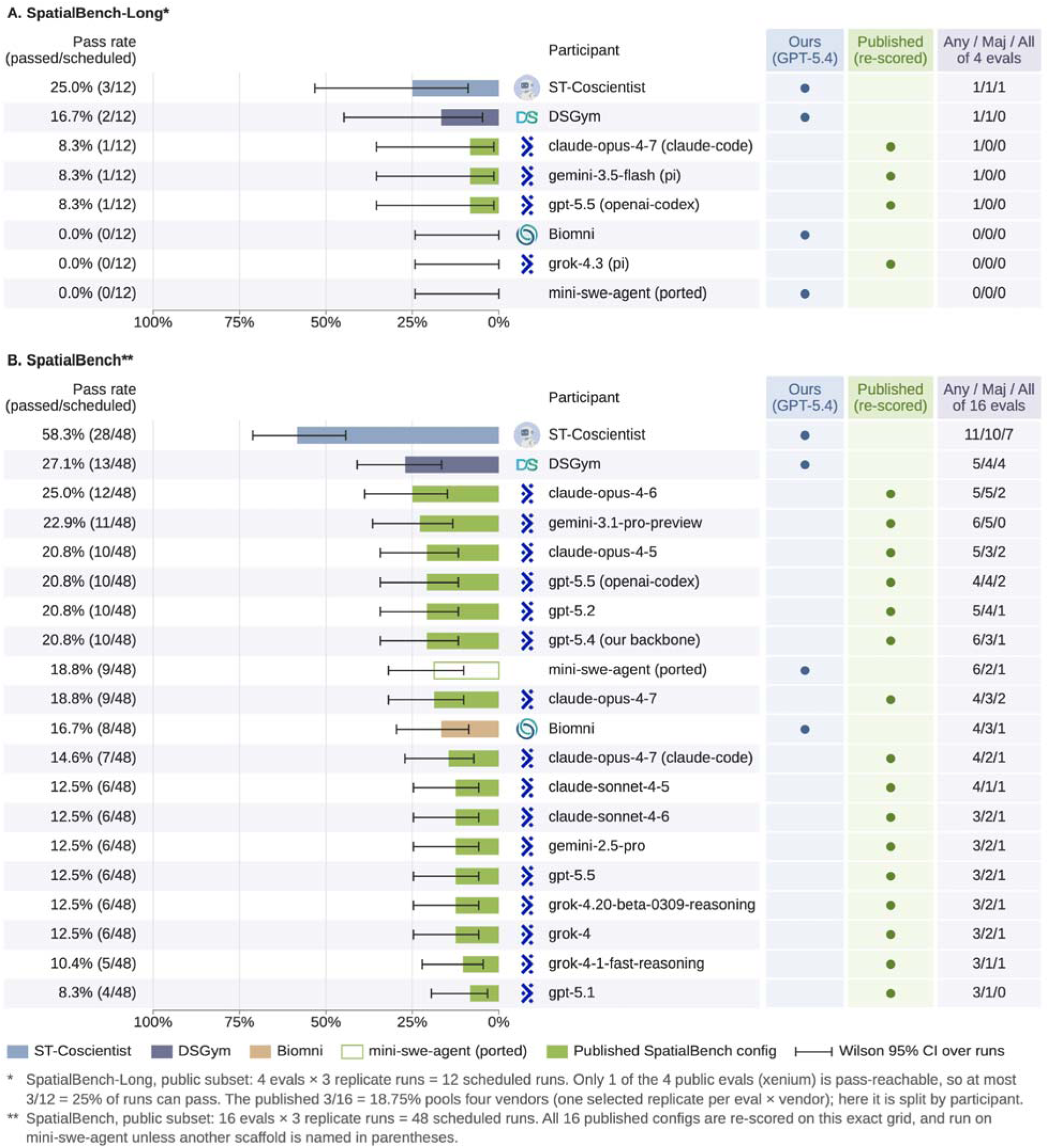
Evaluation on public subsets of SpatialBench and SpatialBench-Long. A, SpatialBench-Long: four evaluations with three scheduled runs each (12 runs). B, SpatialBench: 16 evaluations with three scheduled runs each (48 runs). Bars show run-level pass rates with nominal Wilson 95% confidence intervals, which do not account for dependence among runs of the same evaluation. Markers distinguish GPT-5.4 runs from published configurations re-scored on the same subsets. Any/Maj/All indicates evaluations passed in at least one, a majority or all three runs. The SpatialBench-Long scoring setup marks only one public evaluation as pass-reachable, corresponding to a maximum of 3/12 successful runs.

**Figure 4.**
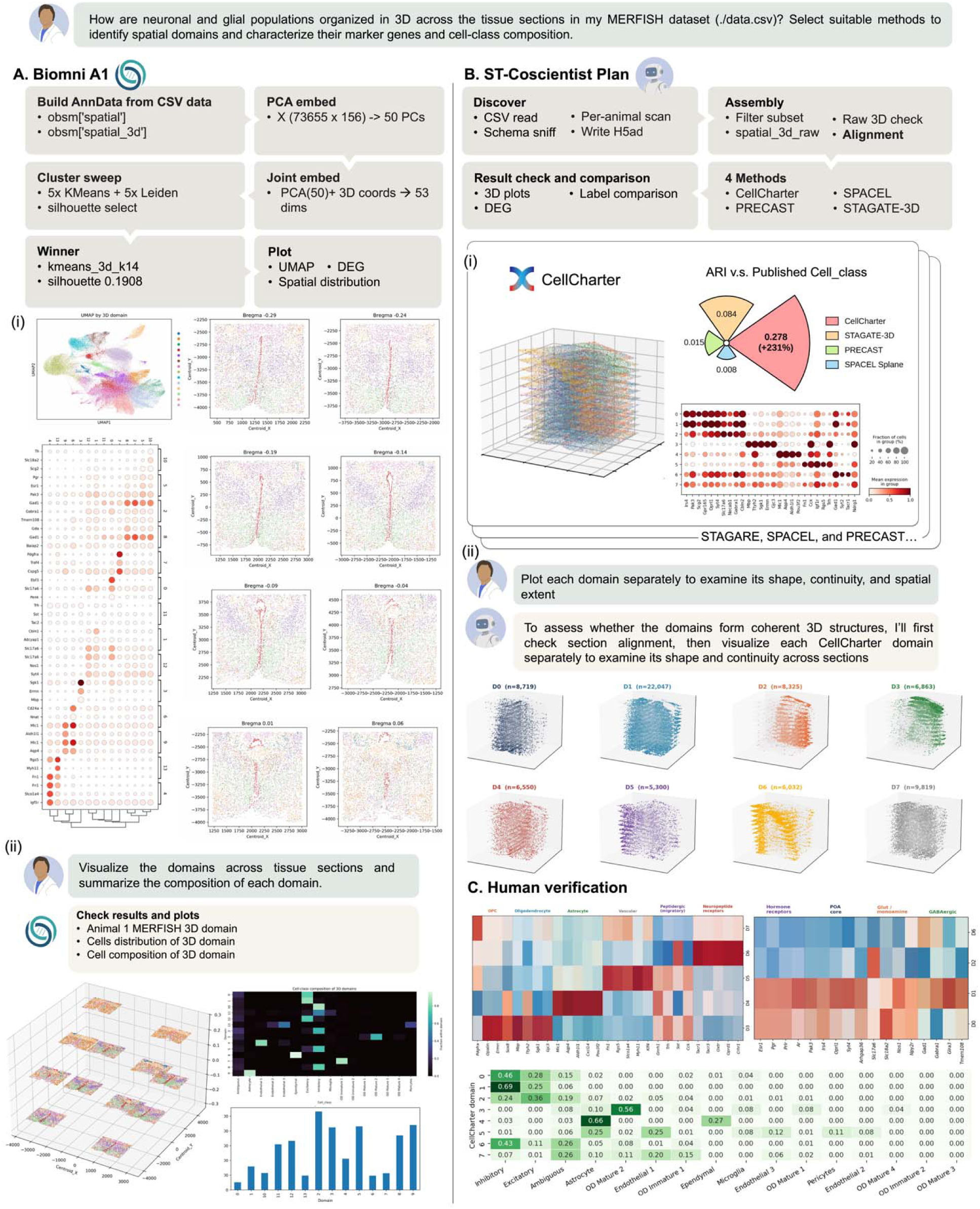
Workflow assembly for user-provided 3D MERFISH data. Top, the user request. A, Biomni A1 constructs an AnnData object, reduces expression to 50 principal components, concatenates these with three spatial coordinates, and compares K-means and Leiden clustering results. It selects a 14-cluster K-means result using the silhouette score and returns embeddings, spatial distributions and marker plots (i). A follow-up request produces 3D and cell-composition views (ii). B, ST-Coscientist performs schema discovery, per-animal scanning, H5AD preparation, coordinate checking and alignment, then compares CellCharter, PRECAST, SPACEL and STAGATE-3D. The displayed CellCharter result includes a 3D view, marker expression and ARI against published cell-class labels (i). A follow-up request generates separate 3D views of the eight CellCharter domains (ii). C, Marker-expression and cell-composition summaries used for human review. Additional method comparisons are provided in Supplementary Fig. 1.

**Figure 5.**
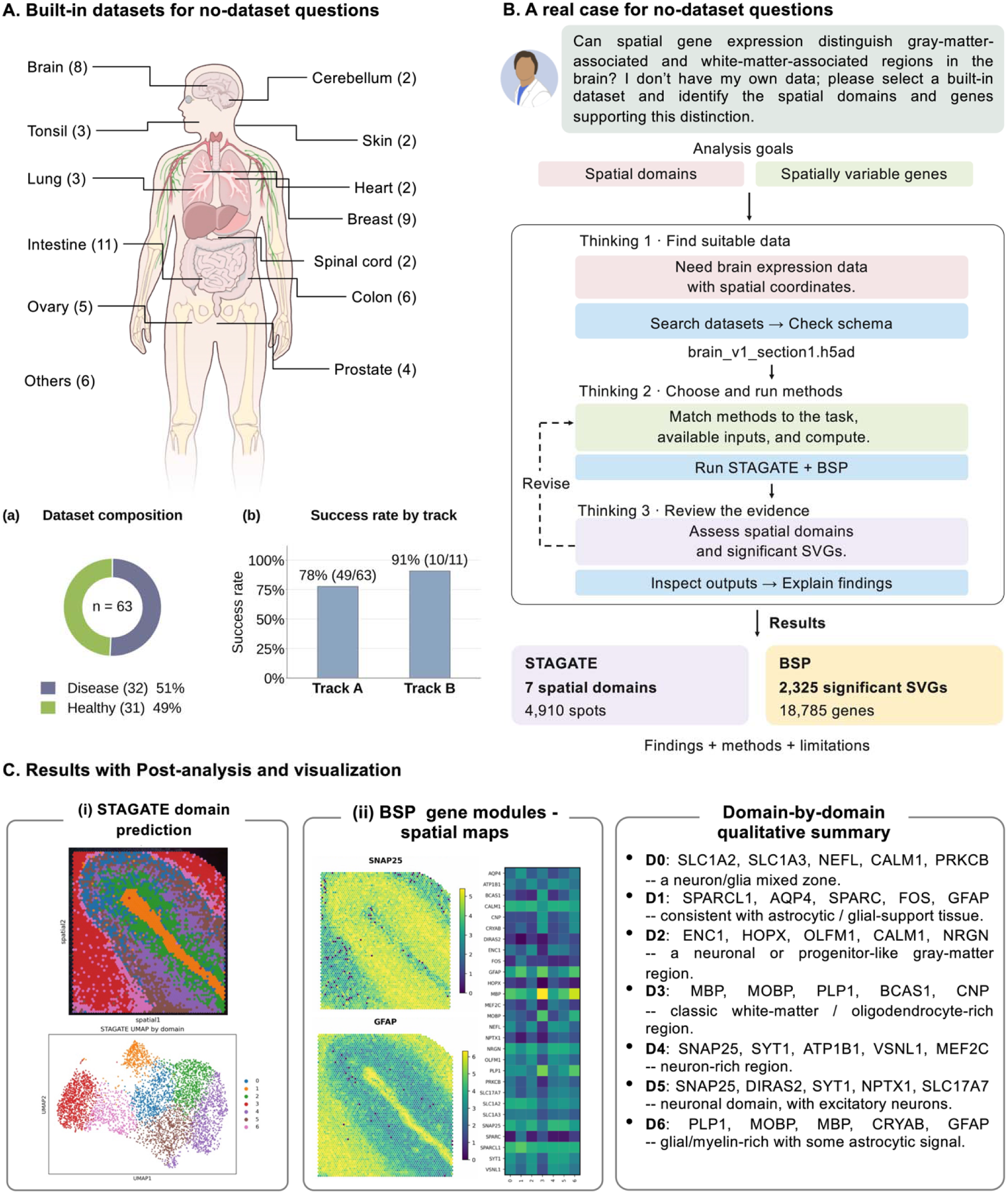
Routing a question posed without data through the built-in dataset library. A, Tissue coverage of the reference library, with dataset counts shown on the body map. Insets show the composition of the 63 disease- or healthy-annotated datasets (a) and retrieval accuracy for single-dataset requests (49/63) and disease–healthy comparisons (10/11) (b). B, For a brain-tissue request without supplied data, ST-Coscientist searches the library, selects brain_v1_section1.h5ad, checks its schema, matches methods to the available inputs and compute, and runs STAGATE and BSP. C, The workflow returns seven STAGATE domains across 4,910 spots, spatial maps and expression summaries for SVG analysis (2,325 significant SVGs among 18,785 genes), and a qualitative domain-by-domain interpretation.

**Figure 6.**
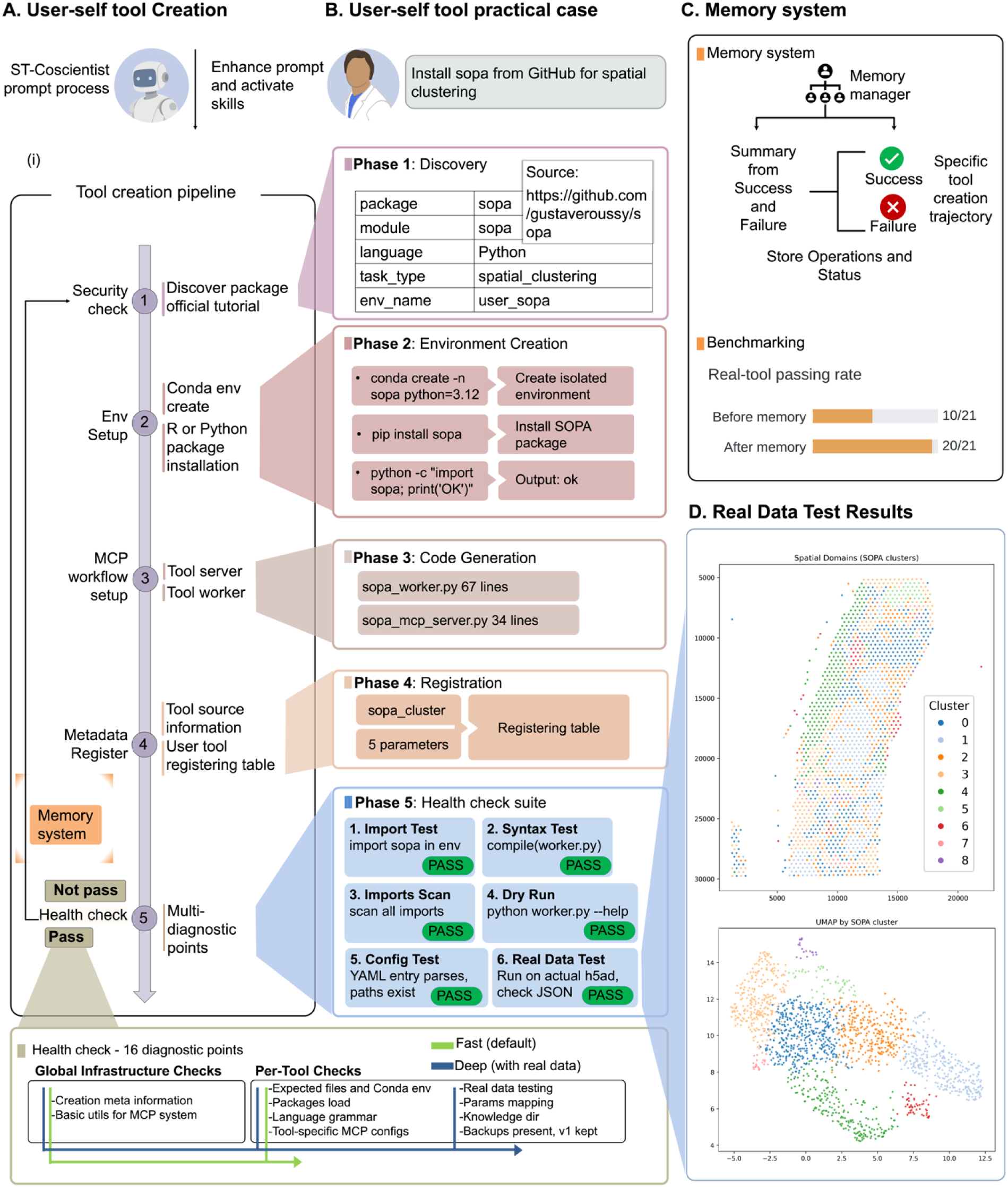
Integration and validation of additional tools through procedural memory. A, Tool-creation pipeline: security check and package discovery, environment setup, MCP worker and server generation, metadata registration, and a 16-point health-check suite (fast infrastructure and per-tool checks by default; deep checks with real data); outcomes are recorded in the memory system. B, Installation of SOPA from GitHub as a spatial-clustering tool: discovery of package metadata, creation of an isolated conda environment, generation of a 67-line worker and a 34-line server, registration of sopa_cluster with five parameters, and six passed checks (import, syntax, dependency scan, dry run, configuration and real data). C, Procedural memory stores summaries and trajectories of successful and failed tool-creation attempts; across 21 GitHub tools, the real-data pass rate rose from 10/21 without memory to 20/21 with memory. D, Real-data test output of the created SOPA tool: spatial-domain map and low-dimensional embedding.

### SpatialOmicsLab extends expert-level performance from execution to analytical choice

SpatialOmicsLab defines 4,614 task specifications spanning 12 task families and three data modalities: 4,004 sequencing-based, 183 image-based and 427 combined sequencing-and-image specifications (Fig. 2A and Supplementary Tables 3 and 4). Within the sequencing-based collection, data preparation and reconstruction encompass quality control and preprocessing (252 specifications), cell segmentation (63), multi-slice alignment (630) and super-resolution (69). Analyses of tissue organization and molecular variation include spatial clustering (948), SVG detection (822) and differential expression (63). Cell-type deconvolution, niche analysis and cell–cell communication contribute 378 specifications each, while single-cell-to-spatial mapping (15) and multi-omics integration (8) connect complementary data sources. Image-based analysis adds 183 cell-segmentation specifications, while combined sequencing-and-image analyses contribute 183 for spatial clustering, 122 for multi-slice alignment and 122 for super-resolution. Each specification connects a research request to a registered analysis pipeline, defining the methods, input requirements and output checks appropriate to the data modality. This organization gives agents a structured basis for assembling workflows around biological questions. We evaluated analytical performance using a separate collection of 103 benchmark tasks with reference annotations (Supplementary Tables 5 and 6).

The benchmarks tested recovery of spatial domains, detection of spatially variable genes and estimation of cell-type composition. Performance was quantified by adjusted Rand index (ARI) against annotated domains, F1 score against reference gene sets and mean Pearson correlation with annotation-derived reference profiles, respectively. A run was scored only after data preparation and method execution yielded a valid prediction that could be aligned with the reference. Runs without an evaluable prediction were recorded as failures, including those whose code completed without error (Methods).

We then compared ST-Coscientist with expert-led workflows using the same methods on the same data and scoring their outputs identically (Fig. 2B). The comparison covered 13 clustering methods on Visium human dorsolateral prefrontal cortex (DLPFC)^62^, seven deconvolution methods on Visium human lymph node with four reference cell types^25^, and eight SVG-detection methods on a Visium mouse brain coronal section with 458 reference SVGs^63^. Expert workflows followed each method’s original documentation and scripts. Within SpatialOmicsLab, ST-Coscientist achieved scores equal to or higher than those of expert workflows across all 28 comparisons. It outperformed expert implementations for methods including stLearn and DeepST in clustering, CARD and SpatialPrompt in deconvolution, and SVGbit and SOMDE in SVG detection.

We next evaluated ST-Coscientist across six benchmark settings in which it selected methods and assembled workflows within SpatialOmicsLab. These settings covered spatial clustering on MERFISH mouse preoptic hypothalamus^64^ and Visium human DLPFC^62^; SVG detection on a Visium mouse brain coronal section^63^; and cell-type deconvolution on Visium human lymph node^25^, Slide-seqV2 human right coronary artery and Slide-seqV2 human aorta^65^. We compared ST-Coscientist with SpatialBench^53^, Biomni^2^ and DSGym^66^. The performance of ST-Coscientist’s backbone language model is shown as a reference (Fig. 2C).

ST-Coscientist outperformed all three comparison systems across the six settings and exceeded the corresponding expert reference scores shown in Fig. 2C. On Visium DLPFC, for example, ST-Coscientist used stLearn to achieve an ARI of 0.509, exceeding the expert reference of 0.487 obtained with BASS. Clustering on MERFISH yielded an ARI of 0.398 versus 0.361 for the expert reference, while SVG detection achieved an F1 score of 0.386 versus 0.341. Mean Pearson correlations for deconvolution were 0.530 versus 0.509 in lymph node, 0.639 versus 0.538 in coronary artery, and 0.546 versus 0.454 in aorta. Within SpatialOmicsLab, ST-Coscientist thus combined expert-level application of established methods with effective analytical choices across measurement platforms.

### SpatialOmicsLab supports long-horizon biological inference

We evaluated ST-Coscientist on SpatialBench^53^ and SpatialBench-Long^60^. SpatialBench tests biological results at defined stages of spatial analysis, whereas SpatialBench-Long asks agents to recover scientific conclusions from raw or near-raw measurements without a prescribed workflow. Within SpatialOmicsLab, ST-Coscientist achieved a SpatialBench pass rate of 58.3%, compared with 27.1% for DSGym^66^ and 16.7% for Biomni^2^, all using GPT-5.4 (Fig. 3B).

The Xenium organoid task in SpatialBench-Long asks how glioblastoma-associated perturbations alter spatial tissue organization. Using matched single-cell and spatial transcriptomic data, agents must identify three niche changes associated with architectural disruption across wild-type and two engineered glioblastoma organoid models, and rank niche abundance across conditions. Answering this question requires linking cell identity to spatial organization while preserving the experimental comparisons that give those patterns biological meaning. ST-Coscientist passed this evaluation in all three runs (Fig. 3A), providing evidence that SpatialOmicsLab supports agent-led investigation in which molecular, spatial and experimental evidence must be brought together to address a biological question.

### SpatialOmicsLab supports adaptive workflows for open-ended biological investigation

We next examined how ST-Coscientist used SpatialOmicsLab to assemble an analysis around the spatial structure of user-provided data. We supplied a raw CSV file containing single-cell-resolution MERFISH measurements from multiple tissue sections of the mouse preoptic hypothalamus^64^. The request was to perform a three-dimensional spatial transcriptomics analysis, leaving method selection and workflow design to the agent. Biomni A1^2^ received the same data and request, allowing us to compare how the two agents recognized analytical prerequisites and organized data preparation, spatial modelling and interpretation (Fig. 4).

Biomni A1 approached three-dimensional analysis as clustering in a combined molecular and spatial feature space (Fig. 4A). It constructed an AnnData object from the 73,655-cell by 156-gene matrix, reduced expression to 50 principal components and appended the three raw spatial coordinates. Within this 53-dimensional representation, it evaluated five K-means and five Leiden solutions and selected a 14-cluster K-means result with a silhouette score of 0.191, followed by UMAP, spatial-distribution and differential-expression plots. The workflow included no explicit section alignment, which can establish a shared coordinate frame for interpreting spatial relationships across tissue sections^67^.

Within SpatialOmicsLab, ST-Coscientist treated spatial correspondence between sections as a prerequisite for domain inference (Fig. 4B). It inspected the CSV schema and the organization of sections by animal, generated standardized H5AD files while retaining the original coordinates in a dedicated spatial_3d_raw field, and checked the coordinate structure before aligning the sections. It then applied four registered spatial-domain methods—CellCharter^68^, PRECAST^69^, SPACEL^70^ and STAGATE-3D^71^—to the aligned data, generating three-dimensional domain maps, differential-expression summaries, domain-composition heatmaps and comparisons with published annotations.

CellCharter showed the strongest agreement with the published cell-class annotations (ARI 0.278; normalized mutual information (NMI) 0.344) and neuron-cluster annotations (NMI 0.232; Supplementary Fig. 1). As cell identities and spatial domains capture different aspects of tissue organization, these scores guided the selection of CellCharter for expert review. In response to a follow-up request, ST-Coscientist rendered each of its eight domains separately in three dimensions, allowing their spatial organization to be examined alongside molecular signatures (Fig. 4B, bottom).

Expert review of the domain maps, marker summaries and differential-expression results supported the interpretation of the eight inferred domains as tissue regions with distinct cellular compositions and molecular profiles (Fig. 4C). D0 (8,719 cells) contained both inhibitory and excitatory neurons and showed monoamine-associated and hormone-receptor signatures, including *Slc18a2*, *Esr1*, *Pgr* and *Prlr*^64^. D2 (8,325 cells) showed the strongest *Slc17a6* expression and the highest proportion of excitatory neurons among the domains, consistent with a glutamatergic-associated profile^64^. D1 (22,047 cells) was enriched in inhibitory neurons and expressed *Gad1* and the androgen receptor *Ar*, together with the receptor genes *Gabra1* and *Glra3*. D6 (6,032 cells) showed enriched expression of Gad1, Syt2 and Gabra1, and 43% of its cells were inhibitory neurons. It also shower a neuropeptide-receptor-rich profile including *Tacr1*, *Tacr3*, *Oxtr*, *Oprd1* and *Crhr1*^64^. D3 (6,863 cells) showed an oligodendrocyte and myelin-associated profile marked by *Mbp*, *Opalin*, *Ermn* and *Sox8*^72,73^. D4 (6,550 cells) showed astrocyte-associated expression of *Aqp4*, *Aldh1l1* and *Mlc1* and comprised predominantly astrocytes (66%) and ependymal cells (27%)^74^. D5 (5,300 cells) showed vascular-associated expression of *Myh11*, *Klf4*, *Fn1*, *Rgs5* and *Slco1a4* and contained endothelial cells and pericytes^75,76^. D7 (9,819 cells) combined oligodendrocyte precursor cell-associated *Pdgfra* expression with immature oligodendrocyte and endothelial cell annotations, supporting a mixed glial–vascular interpretation^64,72^. Three-dimensional views showed distinct distributions of domains with neuronal (D0, D1, D2 and D6) and non-neuronal (D3, D4, D5 and D7) signatures across tissue sections (Supplementary Figs. 2 and 3).

SpatialOmicsLab thus supported an investigation that evolved from workflow design to biological review. ST-Coscientist adapted the analysis to the structure of the data and refined its outputs through further interaction, bringing molecular and spatial evidence together for expert interpretation.

### SpatialOmicsLab connects research questions to suitable data for biological interpretation

We next asked whether ST-Coscientist could initiate an investigation from a biological question without user-provided data. SpatialOmicsLab supports this mode of inquiry through a library of 63 curated 10x Visium datasets covering 12 human tissues and additional sites, comprising 32 disease and 31 healthy datasets (Fig. 5A). In an evaluation of data selection, ST-Coscientist retrieved the correct dataset for 49 of 63 requests requiring a single dataset (78%) and the correct pair for eight of nine requests comparing disease and healthy samples (91%).

For a request to identify spatial domains and spatially variable genes in brain tissue without supplied data, ST-Coscientist retrieved brain_v1_section1.h5ad from the library (Fig. 5B). It inspected the coordinates, count matrix, image availability and uniqueness of gene names to establish the inputs available for analysis. It considered candidate-method benchmark performance together with compatibility with the available data and computing resources. For spatial-domain detection, it retained methods that did not require a GPU, histology image or raw 10x files; for SVG detection, methods had to operate on H5AD-native counts without a single-cell reference. ST-Coscientist selected STAGATE^71^ and BSP^77^ as the highest-ranked compatible methods for the two tasks, respectively.

Spatial-domain and SVG analyses provided complementary views of neuronal and glial organization in the retrieved brain section (Fig. 5C). The workflow identified seven spatial domains across 4,910 spots and 2,325 significant SVGs among 18,785 tested genes. ST-Coscientist returned domain maps, UMAP embeddings, SVG spatial maps, marker heatmaps and summaries of individual domains. Domains 3 and 6 expressed the oligodendrocyte- and myelin-associated genes *MBP*, *MOBP* and *PLP1*^62,72^. Domain 1 showed astrocyte-associated expression of *AQP4*, *GFAP* and *SPARCL1*^74^. Domains 2, 4 and 5 showed neuronal-associated profiles, with Domains 4 and 5 expressing *SNAP25* and *SYT1*, and Domain 5 additionally expressing the glutamatergic marker *SLC17A7*^62,78^. Domain 0 combined astrocytic glutamate-transport signatures (*SLC1A2*, *SLC1A3*) with the neuronal marker *NEFL*, consistent with a mixed neuronal–glial profile^79^. The distinct spatial distributions of *GFAP* and *SNAP25* were consistent with regional differences in astrocytic and neuronal expression^62,74^.

By linking the reference library to data diagnosis and analysis, SpatialOmicsLab supported investigation from a research question without user-supplied data. ST-Coscientist selected the measurements and methods, then assembled the resulting spatial and molecular evidence into a biological interpretation.

### SpatialOmicsLab expands its analytical repertoire through accumulated integration experience

As the spatial-transcriptomics ecosystem evolves^21^, research questions may require methods beyond those already available in the environment. Incorporating such methods requires explicit inputs and outputs, isolated software dependencies and validation on real data. SpatialOmicsLab supports this expansion through a structured integration process (Fig. 6A). Given a source repository and a description of the intended analysis, ST-Coscientist identifies the package, programming language and task type. The package is installed in an isolated Python or R environment, and the agent generates the MCP worker and server, defines the callable interface and registers task metadata. A 16-point diagnostic suite evaluates the resulting tool: fast checks assess infrastructure, required files, imports, syntax and configuration, whereas deep checks test execution on real data, parameter mapping and backup completeness. These checks assess whether an integrated method is ready for subsequent analysis.

ST-Coscientist retrieved SOPA^80^ from GitHub and integrated it as a callable spatial-clustering tool within SpatialOmicsLab (Fig. 6B). It installed the Python package in a dedicated conda environment, generated a 67-line worker and a 34-line server, and registered the sopa_cluster action with five parameters. All six displayed checks passed: package import, syntax, dependency scanning, dry-run execution, configuration parsing and execution on real data. The registered action processed an H5AD dataset and returned spatial-domain assignments and a low-dimensional embedding (Fig. 6D), demonstrating that the added method could execute through the environment’s standard interface.

Procedural memory enabled ST-Coscientist to reuse integration experience when adding further methods. Across 21 methods, the number passing deep validation on real data increased from 10 without memory to 20 with memory (47.6% to 95.2%), without retraining the language model (Fig. 6C and Supplementary Table 7). SpatialOmicsLab retains successful and failed integration trajectories, including encountered errors, attempted repairs and validation outcomes. ST-Coscientist uses prior integration experience to guide setup and repair when similar problems recur, including dependency conflicts, changes in package interfaces, metadata errors, incompatible data structures, hardware assumptions and unexpected output locations. Validation is repeated for each newly integrated or repaired tool.

The analytical repertoire can also be revised as methods and research needs change. User-added tools can be modified, committed, rolled back, deactivated, restored or deleted through the same lifecycle with validation checks (Supplementary Fig. 4). SpatialOmicsLab thus retains both callable methods and reusable integration experience, providing a basis for agents to extend their analytical capabilities as the field evolves.

## Discussion

SpatialOmicsLab addresses the gap between the availability of spatial-omics methods and their use in coherent biological investigations. Our findings show how a domain-specific research environment can connect an agent’s scientific reasoning to executable, traceable analysis. Within SpatialOmicsLab, data context and benchmark evidence inform the agent’s method selection, while standardized interfaces, isolated execution and diagnostic feedback support workflow implementation and revision. Scientific choices and biological interpretation remain with the agent; the environment preserves the computational conditions and records needed to examine its results. We provide SpatialOmicsLab as an extensible research resource, with ST-Coscientist demonstrating how an agent can use it to pursue biological questions.

The benchmarks support expert-level performance in both method implementation and analytical choice. ST-Coscientist achieved scores equal to or higher than those of expert-led workflows across 28 analyses with methods and data held fixed. When it selected methods and assembled workflows, it outperformed the three comparison systems across six settings and exceeded the corresponding expert reference scores. On SpatialBench^53^, ST-Coscientist also outperformed DSGym^66^ and Biomni^2^ when all three used GPT-5.4, extending the evidence to independently defined tasks with a common language-model backbone. Process-level evaluation has likewise revealed substantial performance differences across agent execution frameworks, alongside persistent weaknesses in method selection and biological interpretation despite strong grounding in identifiable sources^55^. These findings argue for evaluating AI co-scientists as complete research systems, including how the environment informs analytical choices and supports their execution.

The case studies illustrate how an investigation can adapt both its workflow and its resources to the biological question. In the MERFISH analysis, ST-Coscientist aligned tissue sections before comparing four domain-detection methods, adapting the workflow to the structure of the measurements. A request without supplied data required the agent to retrieve suitable reference measurements, whereas incorporating an unregistered method expanded the available analytical repertoire. Procedural memory made integration experience reusable: across 21 methods, twice as many passed validation on real data when memory was available. This flexibility is relevant to longer investigations, exemplified by the Xenium organoid task, in which cell identity, tissue organization and experimental comparisons must be considered together. SpatialOmicsLab thus provides a basis for pursuing biological questions whose analytical requirements are not fully specified at the outset.

SpatialOmicsLab complements AI systems for hypothesis generation and collaborative research^34,35^ by providing a setting in which proposed explanations of tissue organization can be examined through traceable spatial analyses. The same environment can connect scientific use with the development and evaluation of agents, complementing benchmarks of task performance and analytical reasoning^48,55^. Its 4,614 task specifications define analytical requests and computational requirements, while separate reference-annotated benchmarks support quantitative evaluation of results. Together, these resources support interactive agent training through task-specific evaluation and execution feedback^52^. Extending this approach to spatial proteomics, spatial multi-omics and imaging would require methods, data representations and evaluation criteria suited to those modalities, while preserving the link between scientific questions and executable analyses.

Computational validation assesses defined requirements for inputs, execution and outputs; method suitability and biological interpretation still require expert scrutiny, particularly in translational or clinical settings. The comparisons evaluate complete agent configurations and do not isolate the contribution of individual environment components. Comparisons with expert workflows covered three task families, while external evaluation sampled public tasks from SpatialBench^53^ and SpatialBench-Long^60^. Broader evaluation is needed for workflows operating directly on images, larger independent cohorts and prospective use by researchers. Reference-data selection remains bounded by the size, diversity and annotation quality of the curated library. Methods obtained from public repositories introduce further challenges as code, dependencies and interfaces change, and the current checks address licensing and security considerations only in part. Expanding the reference library, strengthening repository vetting and version tracking, and extending validation on real data are therefore priorities for broadening the environment’s research use.

SpatialOmicsLab provides a traceable link from biological questions to the analyses and evidence used to address them. ST-Coscientist demonstrates how an agent can work within this environment to adapt its approach to available measurements and develop a biological interpretation. By preserving analytical records and reusable integration experience, the environment provides a foundation for spatial transcriptomic investigations whose methods can evolve with the question and whose results remain open to scientific scrutiny.

## Methods

### Construction of the SpatialOmicsLab environment

#### Environment design

SpatialOmicsLab provides the resources and execution support through which AI agents conduct spatial transcriptomic analyses (Fig. 1). The environment comprises analytical tools, reference datasets, data-preparation utilities, execution interfaces, inspection routines and records of analyses.

#### Tool registry

The registry contains 113 callable functions: 103 analytical tools and 10 utilities (Supplementary Tables 1 and 2). These functions are exposed through 88 Model Context Protocol (MCP) servers. Each entry specifies the analytical task, required data, parameter types and defaults, calling interface, execution environment and expected outputs. The collection includes methods for spatial transcriptomics and utilities for data conversion, dataset search and visualization. A software package may expose multiple registered functions.

#### Reference datasets

A library of 63 reference datasets supports investigations initiated without user-supplied data. Its index records sample identifiers, organism, tissue, disease or healthy status, measurement technology, descriptive metadata and data locations. These annotations support retrieval by the biological context of a question. Retrieved data are assessed for compatibility with candidate methods using the same diagnosis and preparation routines as user-supplied inputs.

#### Task specifications

The task library comprises 4,614 specifications organized by analytical task and data modality: sequencing-based, image-based or combined (Supplementary Tables 3 and 4). Each specification links a research request to an analysis pipeline, defining required inputs, compatible methods and output checks. Specifications describe supported analyses and do not count completed experimental runs. The separate collection of 103 benchmark tasks is described below.

#### Data preparation

Inputs may include prepared spatial datasets, raw platform outputs, single-cell reference data, histology images and associated metadata. Diagnostic utilities describe the measurement platform, file format, spatial-coordinate fields and availability of images, reference data and annotations. ST-Coscientist uses this context to identify preparation steps. Available utilities create or update H5AD objects, extract coordinates, inspect count matrices, standardize gene names, map metadata fields to tool parameters and prepare output locations. Compatibility checks identify unmet requirements before method execution. In the benchmark pipeline, unresolved input problems block execution and are returned as structured failures.

#### Execution interfaces

Registered methods are exposed through MCP servers^61^, which dispatch computation to worker processes in separate Python or R environments (Fig. 1C). The server– worker arrangement accommodates methods with conflicting dependencies while defining their inputs and returned results. ST-Coscientist also has access to Python, R and shell execution for data preparation, diagnostics and supporting analyses.

#### Output inspection and standardization

Task- and tool-specific inspection profiles identify prediction files or object fields, distinguish final results from intermediate files, detect malformed outputs and recover results written to unexpected locations. The benchmark pipeline separates output inspection, standardization and evaluation. Standardized predictions comprise observation-level domain labels for clustering, gene-level tables for spatially variable gene detection, and location-by-cell-type proportion matrices for deconvolution.

#### Tool integration and validation

Additional methods can be incorporated by defining their interfaces, dependencies and expected outputs. The environment provides isolated software environments, worker and server templates, registration metadata, validation routines and records of integration attempts. The 16-point diagnostic suite includes shared infrastructure checks for creation metadata and MCP utilities, together with checks of required files, software imports, code syntax, dependencies, dry-run behavior, configuration, parameter mapping and supporting files or backups. Fast checks omit real-data execution by default; deep checks execute the worker and inspect its outputs. Health reports distinguish critical failures from degraded functionality. Experimental validation criteria are defined below.

#### Tool maintenance

Maintenance functions support changes to tool metadata, server code and worker code (Supplementary Fig. 4). The modification workflow inspects the installed tool, checks for conflicts and prepares temporary changes for validation before committing them; rollback restores the preceding version when an update fails. Removal is staged: a tool can be deactivated and moved to a trash state while retaining its files and metadata. It can subsequently be restored or permanently removed together with its environment and registration.

### Workflow planning and adaptation by ST-Coscientist

#### Task interpretation and resource retrieval

ST-Coscientist accepts research requests accompanied by data, biological questions without data, and requests to incorporate additional methods. A question-management step distinguishes conceptual questions from requests requiring computation. For analytical tasks, the agent retrieves relevant tools, datasets, software resources and procedural guidance. Resource retrieval uses the language model to select from descriptions of available resources; it is distinct from the compatibility-based method recommender. When data are not supplied, the agent can search the reference library and inspect candidate datasets before proceeding (Fig. 5).

#### Prompt enhancement

The research request is augmented with data diagnostics, parameter requirements, ranked methods and example calls (Fig. 1B). Optional components provide guidance for hyperparameter optimization, benchmark evaluation and post-analysis visualization. Reusable know-how documents describe procedures such as preparing single-cell references for deconvolution, retrieving datasets, incorporating external methods, interpreting outputs and consulting procedural memory. These resources provide task-specific context for planning; the agent remains responsible for composing the workflow and revising its choices as results become available.

#### Workflow recommendation

The recommender filters candidates whose requirements are incompatible with the available data, including missing spatial coordinates, images, single-cell references or convertible Visium inputs. Compatible methods are ranked using curated priorities and available benchmark scores for the task and measurement platform. Recommendations include supporting rationales and example calls. When no method is specified, the agent is directed to use the highest-ranked compatible candidate unless a data-driven reason supports another choice. An explicitly requested method takes precedence over these defaults.

#### Iterative execution and adaptation

The agent alternates reasoning, execution and observation^31^ (Fig. 1C). Before an analytical step, it uses inspection routines to assess required files, data compatibility, supporting inputs and output locations. It then calls a registered method or executes supporting code, examines returned logs and results, and determines the next action. Execution errors, missing files or fields, and failed checks can prompt data re-preparation, parameter adjustment, output recovery or method substitution. The workflow continues until the agent produces a final response or encounters an unrecoverable failure, or reaches an execution limit.

#### Self-critique

An optional self-critique stage examines the accumulated reasoning, executions and proposed solution when the agent attempts to finish. Feedback is returned to the agent for additional analysis, up to a configured number of critique rounds.

#### Workflow integration and procedural memory

For a method requested from GitHub, ST-Coscientist inspects the source repository, identifies its package, language and analytical purpose, and uses the environment’s facilities to provision dependencies, generate a worker and MCP server, register the interface and inspect validation results (Fig. 6A). When enabled, procedural memory records the source, environment setup, generated files, errors, repairs and outcomes of successful and failed integration attempts. These records document recurring dependency conflicts, package-interface changes, metadata errors, data-structure mismatches, hardware assumptions and unexpected output locations. The agent can retrieve prior experience to guide installation, metadata mapping, data conversion and output detection. Memory provides guidance for subsequent attempts; it does not replace validation or update the language model’s weights.

#### Visualization and reporting

Post-analysis guidance supports task-specific summaries, interpretation and visualization. Outputs include H5AD or CSV files and saved figures, accompanied by conversation and execution records. Reports present the research request, methods and parameters, analytical steps, output locations, diagnostic messages and interpretation notes (Fig. 1D). The implementation supports HTML analysis reports and PDF export of conversation histories.

### Experimental design and comparative evaluation

#### Evaluation design

We evaluated analytical performance, completion of externally defined tasks, adaptation to research requests and integration of additional methods. Experiments used reference-annotated benchmarks, comparisons with expert workflows and scientific agents, analyses of user-provided and retrieved data, and a comparison of tool integration with and without procedural memory. These experiments assessed complete agent configurations; comparisons using a common language model are identified below.

#### Internal benchmark tasks

A collection of 103 reference-annotated tasks supported quantitative evaluation (Supplementary Tables 5 and 6). Each task specified the data, task family, analytical method, required inputs, expected output structure, reference standard and primary metric. The evaluation covered spatial clustering, spatially variable gene (SVG) detection and cell-type deconvolution, which produce domain assignments, gene-level predictions and cell-type composition estimates, respectively.

#### Expert-led analyses

ST-Coscientist and expert analysts applied the same methods to the same data, and their outputs were scored identically (Fig. 2B). The 28 analyses comprised 13 clustering methods on Visium human dorsolateral prefrontal cortex (DLPFC), sample 151673, with seven annotated tissue regions^62^; seven deconvolution methods on Visium human lymph node with four reference cell types^25^; and eight SVG-detection methods on a Visium adult mouse brain coronal section with 458 reference SVGs^63^. Expert workflows followed the methods’ original documentation or established practice.

#### Agent-selected analyses

Six settings allowed ST-Coscientist to select methods and assemble workflows (Fig. 2C). Clustering was evaluated on MERFISH mouse preoptic hypothalamus with eight reference classes^64^ and Visium human DLPFC^62^; SVG detection on the Visium mouse brain section^63^; and deconvolution on Visium human lymph node^25^, Slide-seqV2 human right coronary artery with six reference lineages and human aorta with seven vascular lineages^65^. Comparison systems were SpatialBench^53^, Biomni^2^ and DSGym^66^, which received the same analysis requests and input data. ST-Coscientist’s backbone language model and the corresponding expert workflow for each setting were shown as references.

#### Language-model configurations

Unless otherwise indicated, ST-Coscientist used GPT-5.4. Performance across language models was examined using the five configurations labelled GPT-5, GPT-5.2-codex, GPT-5.3-codex, GPT-5.4 and GPT-5.4-mini (Supplementary Fig. 5). The evaluation covered clustering on DLPFC sample 151673, deconvolution on the human lymph node and SVG detection on the mouse brain section. Analytical scores were accompanied by counts of OUTPUT_INVALID log events for the four configurations other than GPT-5.

#### External benchmarks

SpatialBench^53^ evaluates results at defined stages of spatial analysis, whereas SpatialBench-Long^60^ evaluates scientific conclusions obtained from raw or near-raw measurements without a prescribed workflow. Public subsets comprised 16 SpatialBench tasks and four SpatialBench-Long tasks, each with three scheduled runs per system (48 and 12 runs, respectively; Fig. 3). New runs of ST-Coscientist, DSGym, Biomni and a ported mini-swe-agent used GPT-5.4. Published configurations re-scored on these task sets were presented separately from the new runs. The Xenium organoid task required identifying three niche changes associated with architectural disruption across wild-type and two engineered glioblastoma organoid models and ranking their abundance using matched single-cell and spatial data.

#### Workflow adaptation

A CSV file containing 73,655 cells and 156 genes from multi-section MERFISH measurements of the mouse preoptic hypothalamus^64^ was supplied to ST-Coscientist and Biomni A1^2^ with the same open-ended request for three-dimensional analysis (Fig. 4). ST-Coscientist inspected the schema and organization by animal, generated H5AD files retaining original coordinates in spatial_3d_raw, checked and aligned coordinates, and applied CellCharter^68^, PRECAST^69^, SPACEL^70^ and STAGATE-3D^71^. Outputs included domain maps, differential-expression summaries, domain composition and comparisons with published annotations. Biomni combined 50 expression principal components with three raw coordinates, evaluated five K-means and five Leiden solutions, and selected a 14-cluster K-means solution by silhouette score. Follow-up requests produced additional three-dimensional and cell-composition views; ST-Coscientist rendered its eight CellCharter domains separately. CellCharter results were selected for expert review on the basis of agreement with the published annotations.

#### Reference-data selection

The reference library contained 63 curated Spatial transcriptomics datasets spanning 12 human tissues and additional anatomical sites. These included 60 10x Visium datasets, 2 10x Visium HD datasets, and 1 Xenium dataset. Of these, 32 were annotated as diseased and 31 as healthy (Fig. 5A). Retrieval was evaluated with 63 single-dataset requests and nine requests for disease–healthy comparisons. For a brain-tissue request without supplied data, ST-Coscientist selected brain_v1_section1.h5ad and inspected coordinates, counts, images and gene-name uniqueness (Fig. 5). It considered benchmark rankings together with input and computing requirements, selecting clustering methods that did not require a GPU, histology image or raw 10x files, and SVG methods compatible with H5AD counts without a single-cell reference. It ran STAGATE^71^ and BSP^77^ on 4,910 spots and assessed 18,785 genes for spatial variability.

#### Method integration

The integration example used SOPA^80^, retrieved from GitHub for spatial clustering (Fig. 6B,D). ST-Coscientist identified the package and intended task, installed it in a dedicated conda environment, and generated a 67-line worker and a 34-line server exposing sopa_cluster through five parameters. Six displayed checks assessed package import, syntax, dependencies, dry-run execution, configuration parsing and execution on real data. The real-data test used an H5AD dataset and inspected the returned domain assignments and low-dimensional embedding.

#### Procedural-memory comparison

Tool integration was evaluated across 21 methods with and without procedural memory, without retraining the language model (Fig. 6C and Supplementary Table 7). The memory condition made records of previous integration attempts available to guide setup and repair. The comparison assessed whether the resulting tools passed deep validation on real data.

### Outcome assessment and reproducibility

#### Reference alignment and analytical metrics

Predictions were aligned with the relevant reference annotations before scoring. Clustering was scored by adjusted Rand index (ARI), measuring chance-adjusted agreement between predicted and reference partitions. SVG detection was scored by F1, the harmonic mean of precision and recall for overlap between predicted SVGs and the reference gene set. For deconvolution, the evaluator represents reference annotations as one-hot indicators at each location. Pearson correlation between predicted cell-type profiles and these indicators is calculated across locations and averaged across evaluable reference classes. This measures agreement with annotated spatial distributions rather than recovery of independently measured cell-type fractions. MERFISH domain assignments were compared with published cell-class and neuron-cluster annotations using ARI and normalized mutual information (NMI). The neuron-cluster comparison in Supplementary Fig. 1 covered all 73,655 cells and 70 reference labels, including the category labelled NaN for non-neuronal cells.

#### Evaluable outputs and failures

A benchmark output was evaluable only after data preparation and method execution produced a prediction that could be aligned with the reference and scored. Missing files, malformed outputs, incompatible structures and non-computable metrics were recorded as failures even when execution completed without an error. Analytical scores and completion status were treated as distinct outcomes. The OUTPUT_INVALID panel in Supplementary Fig. 5 reports counts of diagnostic log events rather than the proportion of tasks that failed.

#### External task completion

External benchmarks were summarized by run-level pass rate, using the number of scheduled runs as the denominator: 48 for SpatialBench and 12 for SpatialBench-Long (Fig. 3). Wilson 95% confidence intervals accompanied these proportions. Repeatability was summarized for each evaluation by Any, Maj and All, indicating success in at least one, at least two or all three runs, respectively. New GPT-5.4 runs and re-scored published configurations retained their separate identities. Under the reported SpatialBench-Long scoring setup, only the Xenium evaluation was pass-reachable, limiting attainable passes to three of the 12 scheduled runs.

#### Retrieval and biological interpretation

A reference-data selection was scored as correct when the selected dataset matched the tissue and condition specified in the request. Accuracy was reported separately for single-dataset requests and disease–healthy requests. Biological interpretation of the MERFISH domains used spatial distributions, marker expression, differential-expression summaries and cell composition, followed by expert review.

#### Tool-integration outcomes

The tool-integration experiment used deep validation on real data as its endpoint. Checks examined whether the generated interface could execute the intended method and return the expected structured outputs, assessing computational behavior rather than biological interpretation. Ten of 21 methods passed without memory and 20 of 21 passed with memory (47.6% and 95.2%, respectively). Integration records retained errors, attempted repairs and validation outcomes for inspection and subsequent use.

#### Statistical units and reproducibility

Performance was summarized separately for the 28 analyses with specified methods, six settings allowing agent selection of methods, individual external-benchmark tasks and the 21 methods in the integration experiment. External benchmarks included three scheduled runs per task, summarized as run-level pass rates and task-level repeatability. Cell and spot counts describe the input measurements.

## Reporting summary

Further information on research design is available in the Nature Portfolio Reporting Summary linked to this article.

## Data availability

This study analysed previously published spatial transcriptomic datasets, as described in the Methods and Supplementary Tables 5 and 6. Released benchmark inputs, reference annotations, task prompts, agent trajectories and evaluation records can be accessed at https://huggingface.co/datasets/JioChiang/SpatialOmicsLab. The reference-data index and released benchmark configurations are available at https://github.com/jiangyi01/SpatialOmicsLab.

## Code availability

Source code for SpatialOmicsLab and ST-Coscientist, including the tool registry, execution interfaces, environment specifications and benchmark scripts, is publicly available at https://github.com/jiangyi01/SpatialOmicsLab under the GNU Affero General Public License v3.0 (AGPL-3.0-only). Third-party tools retain their respective licences.

## Acknowledgements

We thank the investigators who made their spatial transcriptomic datasets publicly available and the developers of the open-source methods used in this study.

## Author contributions

Y.J., X.Z., G.X., W.S. and Y.X. conceived and designed the study; developed the methodology and software; curated and analysed the data; conducted the computational experiments; validated and visualized the results; and wrote and revised the manuscript. P.Q., R.W., J.Y. and J.M. conducted the human evaluation. F.W. and B.Y. developed and implemented the deployment infrastructure.

## Additional information

Supplementary information accompanies this manuscript. Correspondence and requests for materials should be addressed to Guanghua Xiao, Wenqi Shi or Yang Xie.

