## Supplementary Materials for "SpatialOmicsLab: an integrated research environment for AI co-scientists in spatial transcriptomics"

Supplementary figures

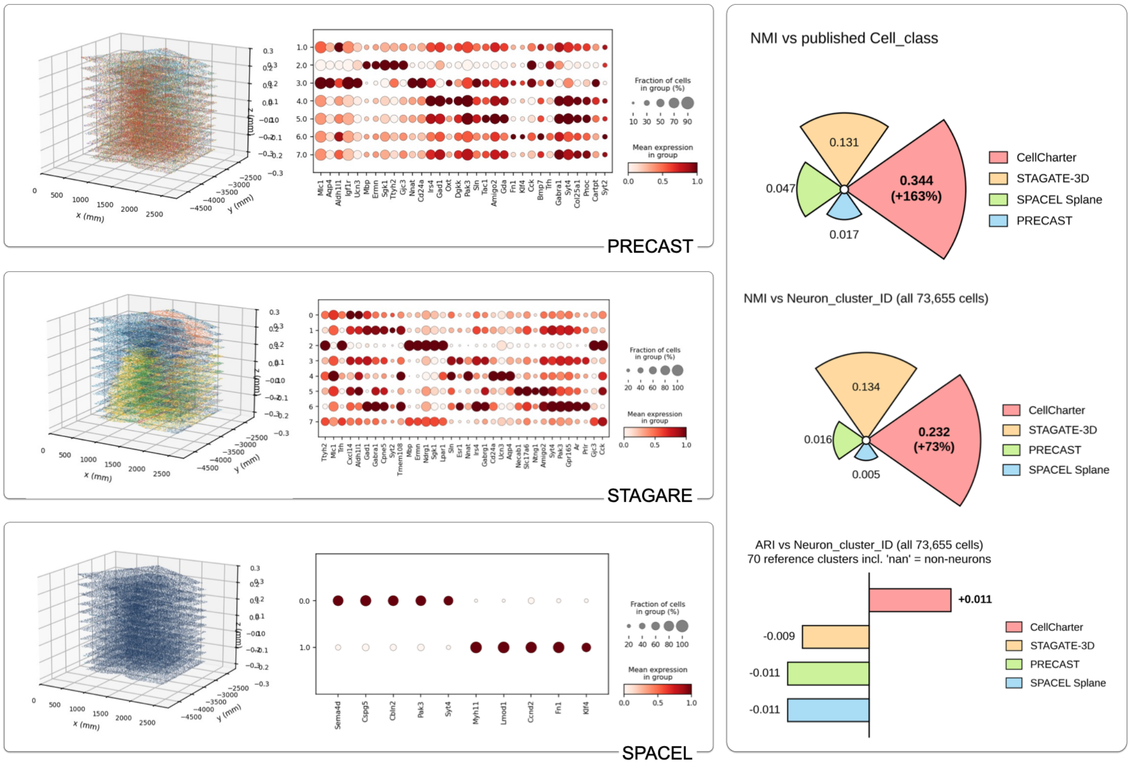


**Supplementary Figure 1 | Comparison of spatial-domain methods on 3D MERFISH data.** Comparison of CellCharter, PRECAST, STAGATE-3D and SPACEL Splane on the mouse preoptic hypothalamus MERFISH dataset. Left, three-dimensional domain maps and representative gene-expression dot plots for PRECAST, STAGATE-3D and SPACEL Splane, from top to bottom. Colours in the spatial maps distinguish inferred domains. Dot size indicates the fraction of cells expressing each gene, and dot colour indicates mean expression within each domain. Right, agreement with published annotations, measured by normalized mutual information (NMI) against cell-class labels (top), NMI against neuron-cluster labels (middle), and adjusted Rand index (ARI) against neuron-cluster labels (bottom).

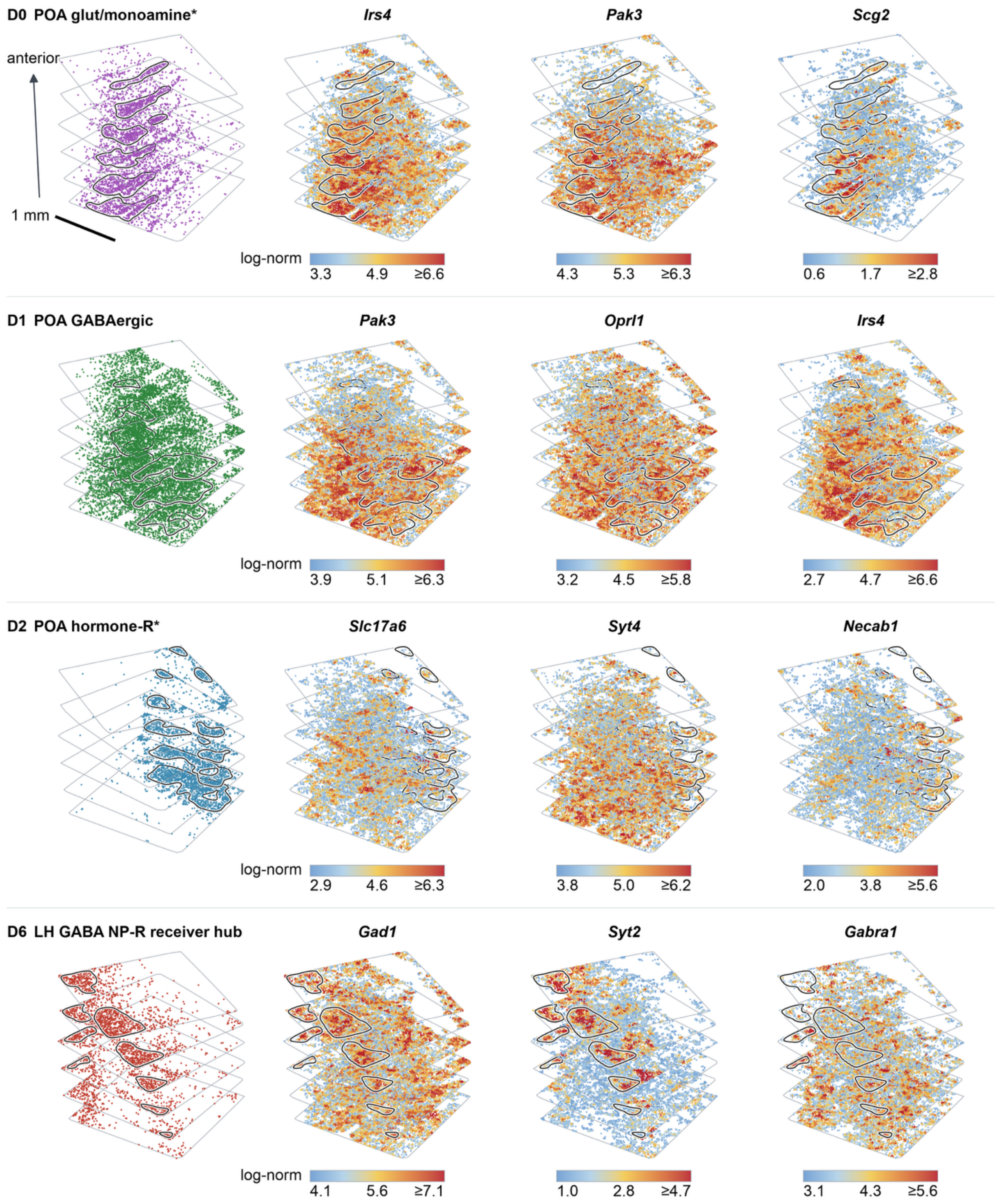


**Supplementary Figure 2 | Neuronal domains in 3D MERFISH data.** Three-dimensional distributions of neuronal domains D0, D1, D2 and D6 inferred by CellCharter in the mouse preoptic hypothalamus MERFISH dataset. Each row shows the spatial distribution of one domain at left and expression maps for three representative genes at right: *Irs4*, *Pak3* and *Scg2* for D0; *Pak3*, *Oprl1* and *Irs4* for D1; *Slc17a6*, *Syt4* and *Necab1* for D2; and *Gad1*, *Syt2* and *Gabra1* for D6. Gene-specific colour scales indicate log-normalized expression, with upper values capped at the indicated limits.

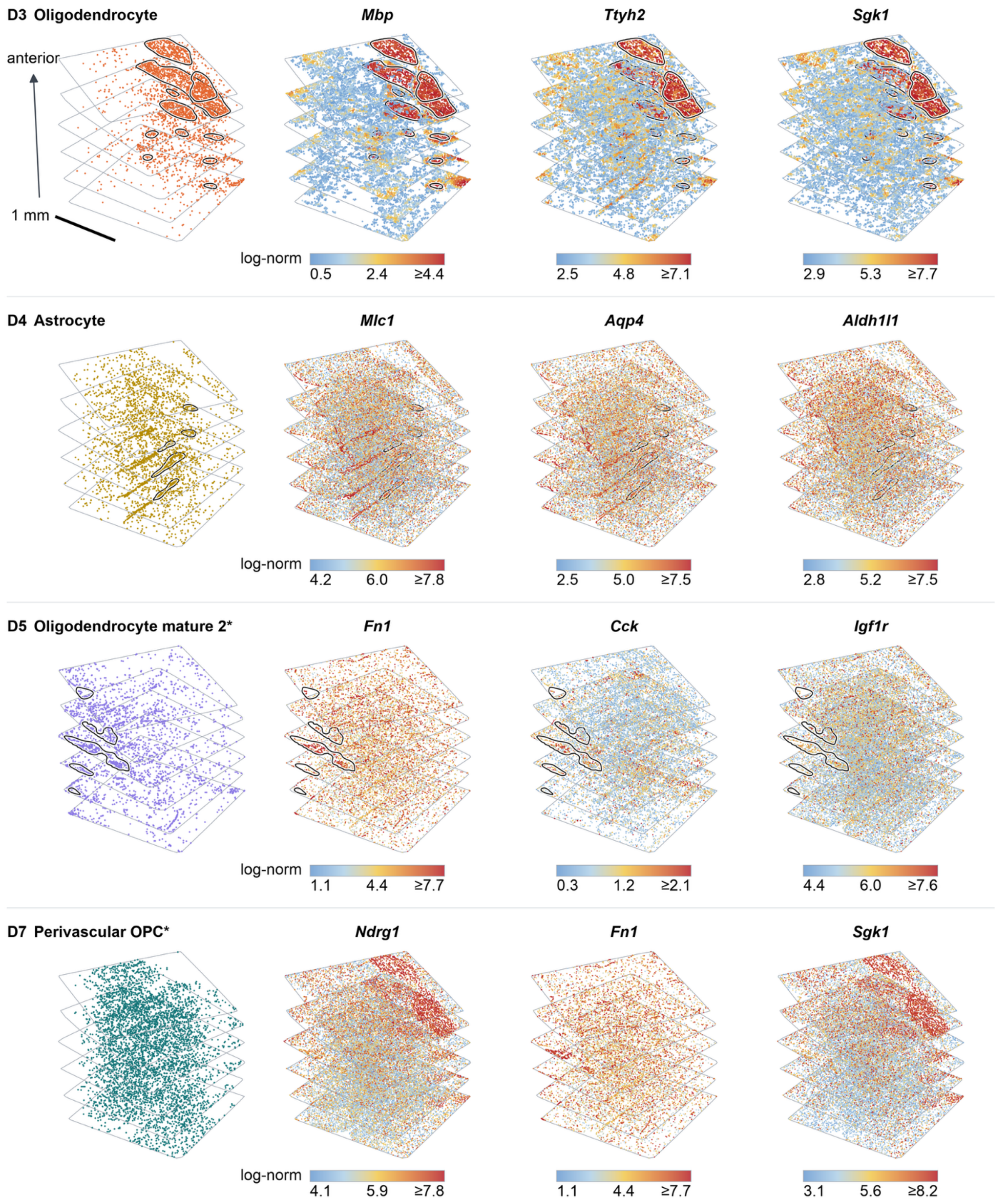


**Supplementary Figure 3 | Non-neuronal domains in 3D MERFISH data.** Three-dimensional distributions of non-neuronal domains D3, D4, D5 and D7 inferred by CellCharter in the mouse preoptic hypothalamus MERFISH dataset, shown alongside expression maps for the representative genes indicated. Spatial distributions, marker expression and cell-class composition support the interpretation of these domains. Domain identifiers correspond to those in Fig. 4. Domain annotations represent putative biological interpretations.

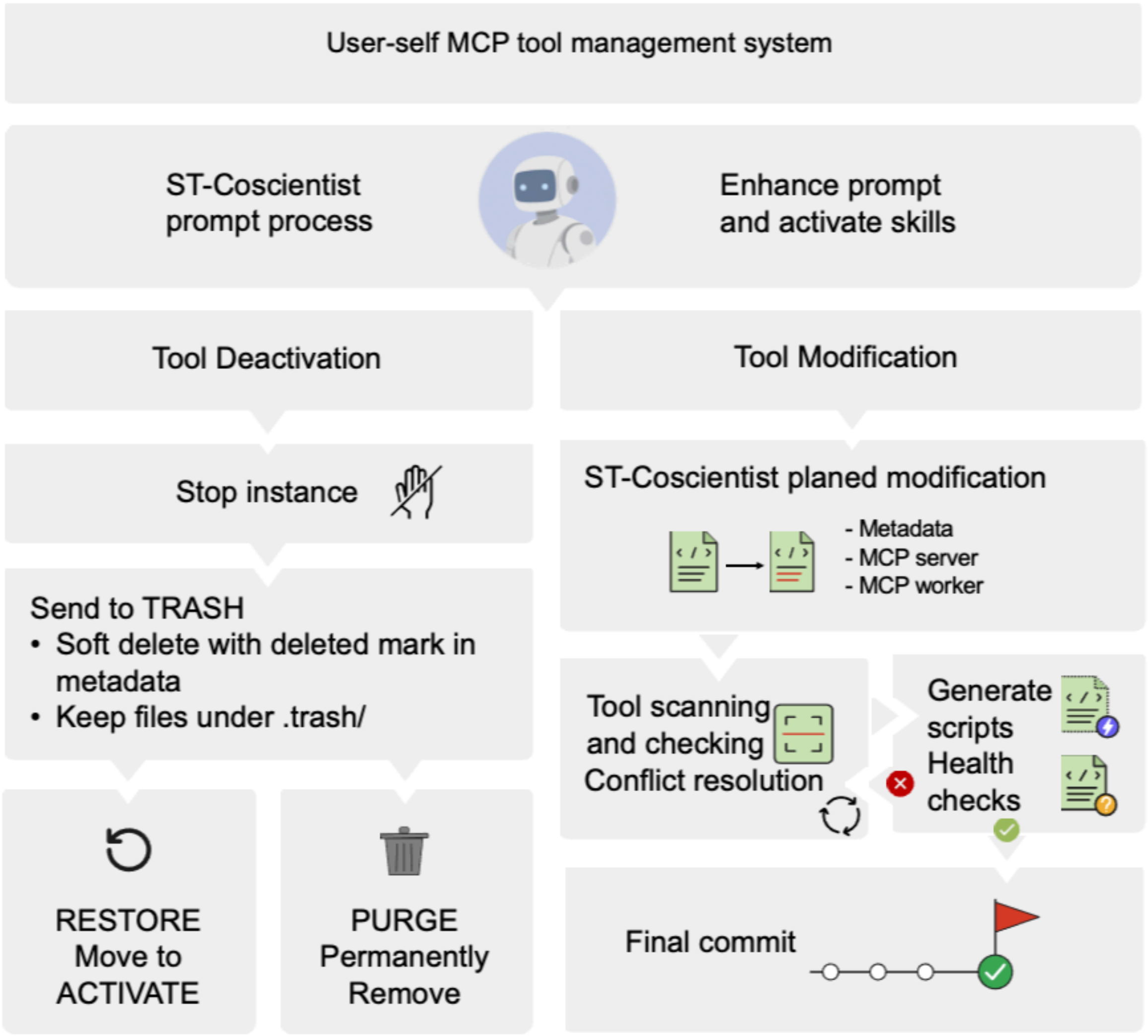


**Supplementary Figure 4 | Management of user-added tools.** ST-Coscientist interprets tool-maintenance requests and invokes procedures for deactivation or modification. Left, deactivation stops the tool instance and moves its files to a recoverable trash state while retaining metadata. The tool can subsequently be restored to active use or permanently removed. Right, modification updates tool metadata, the MCP server or the worker. The workflow inspects the installed tool, resolves conflicts, generates revised scripts and performs health checks before committing the update. Failed checks trigger further revision. MCP, Model Context Protocol.

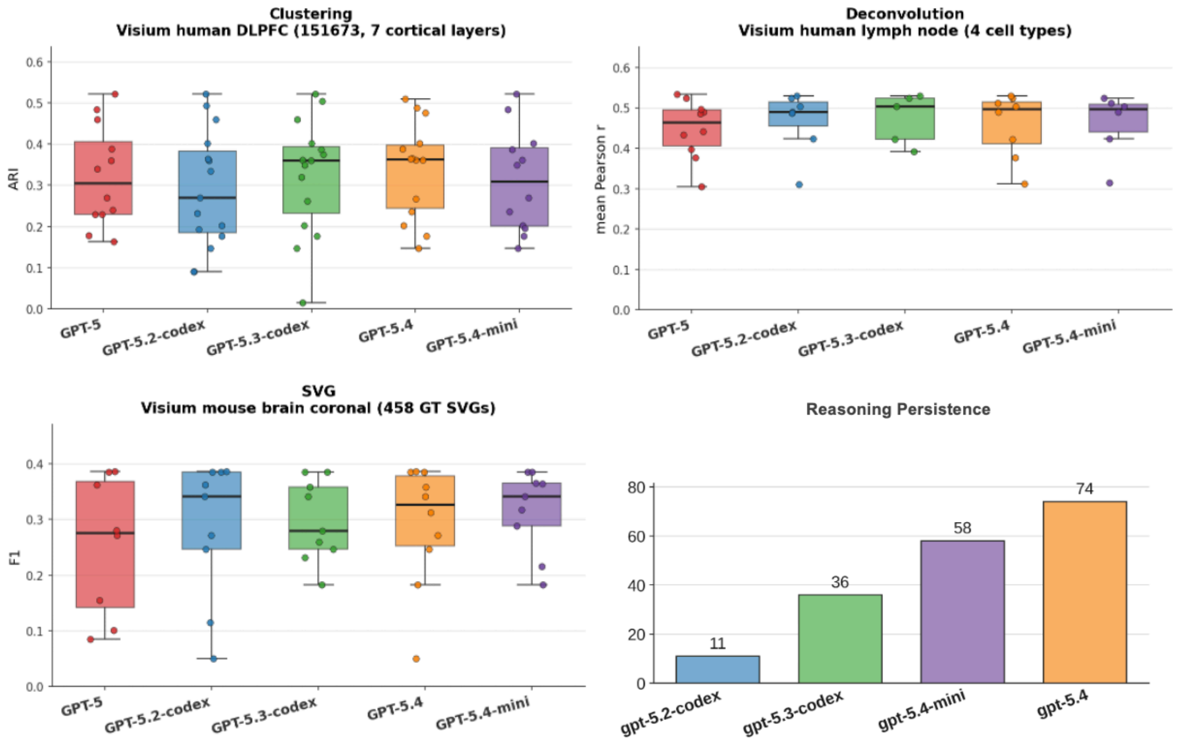


**Supplementary Figure 5 | ST-Coscientist performance across language models.** Comparison of GPT-5, GPT-5.2-codex, GPT-5.3-codex, GPT-5.4 and GPT-5.4-mini within ST-Coscientist. Upper left, spatial clustering of human dorsolateral prefrontal cortex (DLPFC) Visium sample 151673, scored by adjusted Rand index against seven annotated regions. Upper right, human lymph-node deconvolution, scored by mean Pearson correlation across four cell types. Lower left, mouse brain spatially variable gene detection, scored by F1 against 458 reference genes. Points show individual scores; box plots summarize their distributions. Lower right, total reasoning steps used to correct failures during spatial-domain detection on the Visium DLPFC dataset.

**Supplementary Tables**

**Supplementary Table 1 | Registered analytical tools.**

| | Tool | Underlying Package |
|---|---|---|
| 1 | bayestme_deconvolution | BayesTME[1] |
| 2 | bsp_identify_svg | BSP[2] |
| 3 | cellcharter_cluster_spatial_domains | CellCharter[3] |
| 4 | celltrek_spatial_mapping | CellTrek[4] |
| 5 | commot_spatial_communication | COMMOT[5] |
| 6 | deeplinc_cell_interactions | DeepLinc[6] |
| 7 | deeplinc_cell_interactions_csv | DeepLinc[6] |
| 8 | deepst_identify_domains | DeepST[7] |
| 9 | gpsa_align_slices | GPSA[8] |
| 10 | graphst_deconvolution | GraphST[9] |
| 11 | graphst_spatial_clustering | GraphST[9] |
| 12 | hotspot_spatial_modules | Hotspot[10] |
| 13 | istar_full_pipeline | iStar[11] |
| 14 | mist_regions_impute | MIST[12] |
| 15 | mistyr_spatial_modeling | mistyR / MISTy[13] |
| 16 | moscot_run | moscot[14] |
| 17 | ncem_cell_communication | NCEM[15] |
| 18 | neighborseq_interaction_network | Neighbor-seq[16] |
| 19 | novosparc_reconstruct_spatial | novoSpaRc[17] |
| 20 | paste_center_align | PASTE[18] |
| 21 | paste_pairwise_align | PASTE[18] |
| 22 | precast_spatial_clustering | PRECAST[19] |
| 23 | prost_index_svg | PROST[20] |
| 24 | prost_pnn_domains | PROST[20] |
| 25 | redeconve_deconvolution | Redeconve[21] |
| 26 | run_bass | BASS[22] |
| 27 | run_bidcell | BIDCell[23] |
| 28 | run_bulk2space | Bulk2Space[24] |
| 29 | run_card | CARD[25] |
| 30 | run_celina | CELINA[26] |
| 31 | run_cell2location | cell2location[27] |
| 32 | run_celldart | CellDART[28] |
| 33 | run_celloscope | Celloscope[29] |
| 34 | run_cellpie | CellPie[30] |

| | Tool | Underlying Package |
|---|---|---|
| 35 | run_cellpose_segmentation | Cellpose[31] |
| 36 | run_clustermap | ClusterMap[32] |
| 37 | run_cytospace | CytoSPACE[33] |
| 38 | run_deepcell_segmentation | DeepCell / Mesmer[34] |
| 39 | run_destvi | DestVI (scvi-tools)[35] |
| 40 | run_dstg | DSTG[36] |
| 41 | run_gist | GIST[37] |
| 42 | run_iris | IRIS[38] |
| 43 | run_miso | MISO[39] |
| 44 | run_scanpy_spatial_domain | Scanpy[40] / Squidpy[41] |
| 45 | run_scresolve | scResolve[42] |
| 46 | run_sedr | SEDR[43] |
| 47 | run_smart | SMART[44] |
| 48 | run_spacel_scube | SPACEL[45] |
| 49 | run_spacel_splane | SPACEL[45] |
| 50 | run_spatialdecon | SpatialDecon[46] |
| 51 | run_spatialpca | SpatialPCA[47] |
| 52 | run_spatialscope | SpatialScope[48] |
| 53 | run_spicemix | SpiceMix[49] |
| 54 | run_spotclean | SpotClean[50] |
| 55 | run_spotlight | SPOTlight[51] |
| 56 | run_spotsweeper | SpotSweeper[52] |
| 57 | run_stdeconvolve | STdeconvolve[53] |
| 58 | run_stdgcn | STdGCN[54] |
| 59 | run_stplus | stPlus[55] |
| 60 | scdot_map_cells_to_spots | scDOT[56] |
| 61 | seurat_find_markers | Seurat[57] |
| 62 | seurat_integrate_qc_cluster | Seurat[57] |
| 63 | seurat_qc_cluster | Seurat[57] |
| 64 | seurat_spatial_qc_cluster | Seurat[57] |
| 65 | seurat_spatial_variable_features | Seurat[57] |
| 66 | slat_align_slices | SLAT[58] |
| 67 | somde_run | SOMDE[59] |
| 68 | spaceflow_spatial_domains | SpaceFlow[60] |
| 69 | spacet_deconvolution | SpaCET[61] |
| 70 | spacexr_rctd_deconvolution | RCTD[62] |

| | Tool | Underlying Package |
|---|---|---|
| 71 | spagft_identify_svg | SpaGFT[63] |
| 72 | spaotsc_run | SpaOTsc[64] |
| 73 | spark_svg_detection | SPARK[65] |
| 74 | spatialde_run_svg | SpatialDE2[66] |
| 75 | spatialglue_integrate | SpatialGlue[67] |
| 76 | spatialprompt_cluster | SpatialPrompt[68] |
| 77 | spatialprompt_deconvolution | SpatialPrompt[68] |
| 78 | spatrio_align_multiomics | SpaTrio[69] |
| 79 | spiral_align | SPIRAL[70] |
| 80 | spiral_integrate | SPIRAL[70] |
| 81 | spotgf_denoise | SpotGF[71] |
| 82 | spvc_svg_detection | spVC[72] |
| 83 | squidpy_centrality_scores | Squidpy [41] |
| 84 | squidpy_co_occurrence | Squidpy [41] |
| 85 | squidpy_nhood_enrichment | Squidpy [41] |
| 86 | squidpy_ripley | Squidpy [41] |
| 87 | squidpy_spatial_autocorr | Squidpy [41] |
| 88 | squidpy_spatial_neighbors | Squidpy [41] |
| 89 | st_gears_reconstruct_3d | ST-GEARS[73] |
| 90 | stacker_register | ANTsPy[74] / VoxelMorph[75] / SynthMorph[76] (image registration wrapper) |
| 91 | stagate_spatial_domains | STAGATE[77] |
| 92 | stage_run | STAGE[78] |
| 93 | stalign_align_points | STalign[79] |
| 94 | stalign_align_to_image | STalign[79] |
| 95 | starfysh_deconvolution | Starfysh[80] |
| 96 | stlearn_spatial_clustering | stLearn[81] |
| 97 | stride_deconvolution | STRIDE[82] |
| 98 | svca_variance_decomposition | SVCA[83] |
| 99 | svgbit_run | SVGbit[84] |
| 100 | tacco_annotate | TACCO[85] |
| 101 | tangram_map_sc_to_spatial | Tangram[86] |
| 102 | ucdeconvolve_base | UniCell Deconvolve (UCDeconvolve)[87] |
| 103 | xfuse_run | XFuse[88] |

**Supplementary Table 2 | Supporting utilities.**

| | Tool | Underlying Package |
|---|---|---|
| **104** | convert_csv_to_h5ad | in-house utility |
| **105** | convert_h5ad_to_csv | in-house utility |
| **106** | convert_h5ad_to_seurat_rds | in-house utility |
| **107** | convert_seurat_rds_to_h5ad | in-house utility |
| **108** | istar_check_dataset | iStar [dataset check][11] |
| **109** | search_spatial_datasets | in-house utility |
| **110** | seurat_dimplot | Seurat [visualization][57] |
| **111** | seurat_spatial_dimplot | Seurat [visualization][57] |
| **112** | seurat_spatial_feature_plot | Seurat [visualization][57] |
| **113** | xfuse_build_config | XFuse [config builder][88] |

**Supplementary Table 3 | Task coverage by modality and analytical family.**

| Modality | Task | Example methods | Instances |
|---|---|---|---|
| Sequencing-based | Spatial clustering | Scanpy, GraphST, STAGATE, CellCharter, PROST, BASS, PRECAST, SEDR, SpaceFlow, IRIS, SpiceMix, SpatialPCA, SPACEL-Splane, MIST, SpatialPrompt | 948 |
| | Spatially variable gene detection | SpatialDE, SPARK, BSP, SOMDE, SpaGFT, Hotspot, PROST, SVGbit, spVC, SVCA, Squidpy, CELINA, SpotGF | 822 |
| | Multi-slice alignment | PASTE, moscot, ST-GEARS, GPSA, STalign (points), SLAT, SPIRAL, SPACEL-Scube | 630 |
| | Cell-type deconvolution | CARD, cell2location, RCTD, Tangram, DestVI, SPOTlight, TACCO, STRIDE, Starfysh, CellDART, SpatialScope, Bulk2Space-class, STdeconvolve, BayesTME, SpaCET | 378 |
| | Cell-cell communication | COMMOT, SpaOTsc, NCEM, DeepLinc, Neighbor-seq | 378 |
| | Niche analysis | Squidpy (neighbors/nhood/co-occurrence/Ripley/centrality), mistyR | 378 |
| | QC and preprocessing | SpotClean, SpotSweeper, Seurat (QC/integrate) | 252 |
| | Super-resolution (transcript-only) | STAGE, scResolve, stPlus | 69 |
| | Differential expression | Seurat (FindMarkers) | 63 |
| | Cell segmentation (transcript-aware) | ClusterMap | 63 |
| | Single-cell to spatial mapping | novoSpaRc, CellTrek, CytoSPACE, Bulk2Space | 15 |
| | Multi-omics integration | SpatialGlue, SpaTrio | 8 |
| Image-based | Cell segmentation | Cellpose, DeepCell/Mesmer, BIDCell | 183 |
| Sequencing & Image-based | Spatial clustering (H&E-informed) | DeepST, MISO, stLearn | 183 |
| | Multi-slice alignment (image) | STalign (to-image), STACKER | 122 |
| | Super-resolution (histology) | iStar, XFuse | 122 |
| Total | | | 4,614 |

**Supplementary Table 4**

**In additional file: Supplementary Table 4.docx**

**Supplementary Table 5 | Reference-annotated benchmark data.**

| Benchmark dataset | Platform | Task | Tissue (species) |
|---|---|---|---|
| merfish_spatial_domain[89] | MERFISH | Spatial clustering | Preoptic hypothalamus (mouse) |
| visium_dlpfc_domain[90] | Visium | Spatial clustering | Dorsolateral prefrontal cortex, slice 151673 (human) |
| visium_svg[91] | Visium | SVG detection | Brain, coronal; 702 spots (mouse) |
| merfish_svg[92] | MERFISH | SVG detection | Brain receptor-map panel; 649 genes (mouse) |
| merfish_deconvolution[93] | MERFISH | Deconvolution | Primary motor cortex (MOp); 254-gene panel (mouse) |
| visium_deconvolution[27] | Visium | Deconvolution | Lymph node; 4035 spots (human) |
| slideseqv2_aorta_deconvolution[94] | Slide-seqV2 | Deconvolution | Aorta; 6142 beads (human) |
| slideseqv2_rca_deconvolution[94] | Slide-seqV2 | Deconvolution | Right coronary artery; 12744 beads (human) |

**Supplementary Table 6 | Single-cell references for deconvolution.**

| **Single-cell reference** | **Paired benchmark dataset(s)** |
|---|---|
| Mouse MOp scRNA/snRNA-seq taxonomy (subclass_label)[95] | merfish_deconvolution |
| Human secondary-lymphoid-organ scRNA-seq atlas (CellType)[27] | visium_deconvolution |
| Human vascular-tissue scRNA-seq reference, ~113k cells (cell_type)[94] | slideseqv2_aorta + slideseqv2_rca |

**Supplementary Table 7 | Real-data validation of additional tools with and without procedural memory.**

| | Tool name | Task type | Before memory | After memory |
|---|---|---|---|---|
| 1 | BANKSY[96] | Spatial clustering | Yes | Yes |
| 2 | SEAGAL[97] | Gene association | Yes | Yes |
| 3 | SpaGE[98] | Gene imputation | Yes | Yes |
| 4 | SpaVAE[99] | Spatial clustering | Yes | Yes |
| 5 | SpaDecon[100] | Deconvolution | No | Yes |
| 6 | GASTON[101] | Spatial clustering | Yes | Yes |
| 7 | SpaTopic[102] | Spatial clustering | Yes | Yes |
| 8 | semla[103] | Spatial clustering | Yes | Yes |
| 9 | hdWGCNA[104] | Co-expression network | Yes | Yes |
| 10 | SpatialInferCNV[105] | CNV detection | No | Yes |
| 11 | NicheCompass[106] | Cell communication | Yes | Yes |
| 12 | LIANA+[107] | Cell communication | No | Yes |
| 13 | Stereopy[108] | Spatial clustering | No | Yes |
| 14 | Sopa[109] | Segmentation | Yes | Yes |
| 15 | Giotto Suite[110] | Multi-task ecosystem | No | No |
| 16 | Novae[111] | Spatial clustering | No | Yes |
| 17 | Voyager[112] | Gene association | No | Yes |
| 18 | FICTURE[113] | Segmentation | No | Yes |
| 19 | STAMP[114] | Dimension reduction | No | Yes |
| 20 | SpaNorm[115] | Normalization | No | Yes |
| 21 | SPLISOSM[116] | Isoform detection | No | Yes |

Supplementary Information

### 1. Evaluation criteria and statistical metrics

For each evaluated workflow, SpatialOmicsLab first determines whether the run is evaluable. A run is not considered successful only because a process finishes. It must pass the following validation requirements: the required inputs must exist, the selected method must be compatible with the data, the expected output must be produced, the prediction object must be identifiable, and the task-specific metric must be computable. The same evaluation logic is applied to ST-Coscientist, expert hand-run workflows, and general-purpose agent baselines.

Let $b$ denote a benchmark instance, $D_b$ the corresponding dataset, $a$ the execution mode, and $m$ the selected method. The raw output of the workflow is denoted as

$$O_{b,m,a} = f_{m,a}(D_b).$$

The run is considered evaluable only if the output passes validation and can be standardized into the canonical prediction format for the task:

$$E_{b,m,a} = \begin{cases} 1, & \text{if the output is present, valid, standardized, and scoreable,} \\ 0, & \text{otherwise.} \end{cases}$$

Only runs with $E_{b,m,a} = 1$ are assigned a task metric. Failed runs, missing outputs, empty outputs, incompatible schemas, degenerate predictions, and non-computable metrics are recorded as structured failures. For all metrics below, predicted and reference quantities are matched on shared observations, spatial locations, cell types, or gene identifiers before scoring. Clustering and deconvolution metrics are computed with standard Python scientific libraries, including scikit-learn[117] and SciPy[118]. The highly variable gene filter used for SVG benchmarking is computed with Scanpy[40].

#### 1.1 Spatial clustering

Spatial clustering assigns a domain label to each spot or cell. Let

$$Y = \{y_1, \dots, y_n\}$$

be the reference labels over $n$ shared observations, and let

$$\hat{Y} = \{\hat{y}_1, \dots, \hat{y}_n\}$$

be the predicted labels. Let $n_{ij}$ be the number of observations assigned to reference class $i$ and predicted cluster $j$. The row and column sums are

$$a_i = \sum_j n_{ij}, b_j = \sum_i n_{ij}.$$

Writing

$$C(m, 2) = \frac{m(m-1)}{2},$$

the adjusted Rand index is

$$\mathrm{ARI}(Y,\hat{Y}) = \frac{\sum_{ij} C(n_{ij},2) - \frac{\sum_i C(a_i,2)\sum_j C(b_j,2)}{C(n,2)}}{\frac{1}{2}\left[\sum_i C(a_i,2) + \sum_j C(b_j,2)\right] - \frac{\sum_i C(a_i,2)\sum_j C(b_j,2)}{C(n,2)}}.$$

ARI is invariant to label names and corrected for chance. Higher ARI indicates stronger agreement between predicted spatial domains and the reference annotation. ARI is the primary metric for the spatial clustering benchmarks in Fig. 2, including Visium DLPFC and MERFISH clustering tasks.

For completeness, SpatialOmicsLab also reports normalized mutual information, homogeneity, completeness, and V-measure as secondary clustering metrics. With mutual information $I(Y;\hat{Y})$ and entropies $H(Y)$ and $H(\hat{Y})$, normalized mutual information is

$$\mathrm{NMI}(Y,\hat{Y}) = \frac{2I(Y;\hat{Y})}{H(Y) + H(\hat{Y})}.$$

Homogeneity and completeness are computed as

$$h = 1 - \frac{H(Y \mid \hat{Y})}{H(Y)},$$
$$c = 1 - \frac{H(\hat{Y} \mid Y)}{H(\hat{Y})}.$$

The V-measure is

$$V = \frac{2hc}{h + c}.$$

#### 1.2 Cell-type deconvolution

Cell-type deconvolution predicts the spatial distribution of cell types. Let

$$P \in \mathbb{R}^{s \times T}$$

be the predicted spot-by-cell-type matrix over $s$ shared spatial locations and $T$ matched cell types. Let

$$R \in \mathbb{R}^{s \times T}$$

be the corresponding reference matrix. In the implemented evaluator, each row of R is the one-hot encoding of the reference annotation at that location. The score therefore measures spatial agreement with annotation-derived classes rather than recovery of independently measured cell-type fractions. Cell-type names are aligned by exact matching followed by matching after normalizing case, whitespace and separators when needed. The reference matrix is reindexed to the predicted cell-type columns. If the predicted table is transposed, SpatialOmicsLab corrects the orientation during output standardization. Predicted proportions are row-normalized before scoring when required.

For each cell type $t$, Pearson correlation is computed across spatial locations:

$$r_t = \frac{\sum_{s'=1}^{s}(P_{s't} - \bar{P}_t)(R_{s't} - \bar{R}_t)}{\sqrt{\sum_{s'=1}^{s}(P_{s't} - \bar{P}_t)^2}\sqrt{\sum_{s'=1}^{s}(R_{s't} - \bar{R}_t)^2}}.$$

Cell types with near-zero variance in either the predicted or reference vector are excluded:

$$T_{\text{valid}} = \{t: \sigma(P_{\cdot t}) > 10^{-10} \text{ and } \sigma(R_{\cdot t}) > 10^{-10}\}.$$

The primary deconvolution score is the mean Pearson correlation across valid cell types:

$$\text{MeanPearson}(P, R) = \frac{1}{|T_{\text{valid}}|}\sum_{t \in T_{\text{valid}}} r_t .$$

Higher mean Pearson correlation indicates stronger agreement between predicted cell-type profiles and the spatial distribution of the reference annotations. This metric is used for deconvolution benchmarks in Fig. 2, including Visium lymph node and Slide-seqV2 vascular datasets.

SpatialOmicsLab also reports secondary deconvolution metrics. Root-mean-square error is

$$\text{RMSE}(P, R) = \sqrt{\frac{1}{sT}\sum_{s'=1}^{s}\sum_{t=1}^{T}(P_{s't} - R_{s't})^2}.$$

Mean absolute error is

$$\text{MAE}(P, R) = \frac{1}{sT}\sum_{s'=1}^{s}\sum_{t=1}^{T}|P_{s't} - R_{s't}|.$$

For each spatial location, Jensen–Shannon distance is computed between the predicted and reference cell-type distributions. Let

$$M_{s'} = \frac{1}{2}(P_{s'} + R_{s'}).$$

Then

$$\text{JSD}_{s'} = 12\text{KL}(Ps' \,||\, Ms') + 12\text{KL}(Rs' \,||\, Ms')$$

.

Before this calculation, $10^{-12}$ is added to each entry of each row, and both rows are normalized to sum to one; KL uses natural logarithms. The reported value is the mean Jensen–Shannon distance across spatial locations:

$$\text{MeanJSD} = \frac{1}{s}\sum_{s'=1}^{s}\text{JSD}_{s'} .$$

Where predicted cell types are finer than the reference annotation classes, prediction columns are aggregated into the corresponding reference classes using a dataset-specific mapping before scoring.

#### 1.3 Spatially variable gene detection

Spatially variable gene detection returns a set or ranked list of genes predicted to show spatial structure. The predicted SVG set is assembled from the standardized tool output. If $p$-values are reported, genes with $p < 0.05$ are selected; whether the reported values are adjusted depends on the method and output field. If no $p$-value is reported, all returned genes are used subject to the tool-specific top-$N$ rule.

Let

$$S_{\text{pred}}$$

be the predicted SVG set, and let

$$S_{\text{true}}$$

be the benchmark ground-truth SVG set. True positives, false positives, and false negatives are

$$TP = | S_{\text{pred}} \cap S_{\text{true}} |,$$
$$FP = | S_{\text{pred}} \setminus S_{\text{true}} |,$$
$$FN = | S_{\text{true}} \setminus S_{\text{pred}} |.$$

Precision and recall are

$$\text{Precision} = \frac{TP}{TP + FP} = \frac{TP}{| S_{\text{pred}} |},$$
$$\text{Recall} = \frac{TP}{TP + FN} = \frac{TP}{| S_{\text{true}} |}.$$

The primary SVG score is the F1 measure:

$$\text{F1} = \frac{2 \cdot \text{Precision} \cdot \text{Recall}}{\text{Precision} + \text{Recall}}.$$

If no genes are predicted, precision is set to zero. If no benchmark genes are recovered, recall is set to zero. If precision and recall are both zero, F1 is defined as zero. F1 is the primary metric for the Visium mouse brain SVG benchmark in Fig. 2.

To compare methods on a common gene space, SVG inputs are pre-filtered to the top 1,500 highly variable genes using Scanpy with the Seurat flavor. This filter is skipped when the panel contains 1,500 or fewer genes, as in targeted imaging-based platforms. Full-transcriptome significance-test methods whose calibration depends on the complete gene catalog are exempt from this runtime filter. A separate post-hoc analysis intersects each method's predictions with the 1,500-gene set and recomputes F1 within this restricted prediction space.

SpatialOmicsLab also reports Moran's $I$ spatial autocorrelation over predicted genes when applicable. For gene expression vector $x$, spatial weights $w_{ij}$, and $W = \sum_i \sum_j w_{ij}$, Moran's $I$ is

$$I = \frac{n}{W} \frac{\sum_i \sum_j w_{ij} (x_i - \bar{x})(x_j - \bar{x})}{\sum_i (x_i - \bar{x})^2}.$$

Moran's $I$ is used as an auxiliary spatial-structure summary rather than the primary SVG benchmark metric.

#### 1.4 Three-dimensional MERFISH method comparison

For the user-provided 3D MERFISH case, spatial-domain outputs are compared with published annotations using ARI and normalized mutual information. ARI is computed as described in Section 1.1. NMI is used because it measures shared information between two partitions and is invariant to domain-label names.

Let $U$ denote the published annotation and $V$ denote the predicted spatial-domain labels. The mutual information is

$$I(U,V) = \sum_i \sum_j p(i,j) \log \frac{p(i,j)}{p(i)p(j)},$$

where $p(i,j)$ is the joint probability of reference class $i$ and predicted domain $j$, and $p(i)$ and $p(j)$ are the corresponding marginal probabilities. The Shannon entropy of a labeling is

$$H(U) = -\sum_i p(i) \log p(i),$$
$$H(V) = -\sum_j p(j) \log p(j).$$

NMI is computed as

$$\mathrm{NMI}(U,V) = \frac{I(U,V)}{\mathrm{mean}(H(U),H(V))} = \frac{2I(U,V)}{H(U)+H(V)}.$$

In the 3D MERFISH analysis, ARI and NMI compare CellCharter, PRECAST, SPACEL and STAGATE-3D with published cell-class and neuron-cluster annotations (Fig. 4 and Supplementary Fig. 1). Agreement with these annotations was used to select CellCharter for marker-based human review. Because the annotations informed selection, these scores describe this exploratory comparison rather than an independent test of the selected method.

#### 1.5 Baseline model-selection metric

The Biomni A1 baseline in the 3D MERFISH case selects among candidate K-means and Leiden clustering solutions using the silhouette coefficient. For observation $i$, let $a(i)$ be the mean distance from $i$ to other observations in the same cluster. Let $b(i)$ be the lowest mean distance from $i$ to observations in any other cluster. The silhouette value is

$$s(i) = \frac{b(i) - a(i)}{\max(a(i), b(i))}.$$

The overall silhouette score is

$$\text{Silhouette} = \frac{1}{n} \sum_{i=1}^{n} s(i).$$

This score is used only as an internal model-selection criterion for the general-purpose baseline. It is not treated as the biological validation metric for the 3D MERFISH task. ST-Coscientist instead compares spatial-domain methods against published annotations using ARI and NMI, followed by human marker-based review.

#### 1.6 User-added tool validation metric

For runtime tool creation, a GitHub tool is considered validated only if it passes the required health-check suite. Let $T$ be the set of tested user-added tools, and let $C_t$ be the set of required checks for tool $t$. Each check $c \in C_t$ has an indicator

$$I_{t,c} = \begin{cases} 1, & \text{if check } c \text{ passes for tool } t, \\ 0, & \text{otherwise.} \end{cases}$$

The tool-level pass indicator is

$$H_t = \prod_{c \in C_t} I_{t,c}\,.$$

Thus, a tool passes only when all required checks pass. The overall tool pass rate is

$$\text{ToolPassRate} = \frac{1}{|\,T\,|} \sum_{t \in T} H_t\,.$$

Fast-mode checks include metadata validation, expected file presence, conda-environment access, package loading, language syntax, and MCP configuration. Additional diagnostics examine parameter mapping, knowledge-directory integrity and backup consistency. Deep validation includes execution on real data and inspection of its outputs. The memory comparison used deep validation on real data as its endpoint (Fig. 6C and Supplementary Table 7).

#### 1.7 Reference-dataset selection accuracy

For a biological request without user-supplied data, dataset selection is correct when the selected dataset matches the tissue and condition specified in the request. Let $Q$ be the set of evaluated requests. For prompt $q$, define

$$S_q = \begin{cases} 1, & \text{correct selection} \\ 0, & \text{otherwise} \end{cases}.$$

Dataset-selection accuracy is

$$\text{SelectionAccuracy} = \frac{1}{|\,Q\,|} \sum_{q \in Q} S_q\,.$$

For disease–healthy requests, selection is correct only when both selected datasets match the requested tissue and their respective conditions. Accuracy is reported separately for 60 single-dataset requests and nine paired requests (Fig. 5A). Schema validation, method selection and downstream execution are illustrated separately in Fig. 5B,C.

### 2. Task-specification library

The task library contains 4,614 specifications across 12 analytical families and three modalities, represented by 16 modality-specific categories in Supplementary Table 3 (Fig. 2A). Sequencing-based specifications comprise spatial clustering (948), SVG detection (822), multi-slice alignment (630), cell-type deconvolution (378), cell–cell communication (378), niche analysis (378), quality control and preprocessing (252), super-resolution (69), differential expression (63), cell segmentation (63), single-cell-to-spatial mapping (15) and multi-omics integration (8), for a total of 4,004. Image-based cell segmentation contributes 183 specifications. Combined sequencing-and-image analyses comprise spatial clustering (183), multi-slice alignment (122) and super-resolution (122), for a total of 427. A specification defines required inputs, compatible methods and expected outputs; these counts describe supported analyses rather than completed experimental runs. The separate collection of 103 benchmark tasks supports quantitative evaluation.

#### 3. Reference-data retrieval and analysis

SpatialOmicsLab supports biological investigations initiated without user-supplied data through a library of 60 curated 10x Visium datasets: 34 disease and 26 healthy datasets covering 12 human tissues and additional sites (Fig. 5A). ST-Coscientist extracts the tissue, condition and analytical intent from a request, searches the library, diagnoses the selected dataset and ranks methods after assessing compatibility with the available inputs. It selected the correct dataset for 46 of 60 single-dataset requests (77%) and the correct pair for eight of nine disease–healthy requests (89%). In the brain example (Fig. 5B,C), it selected brain_v1_section1.h5ad, inspected coordinates, counts, images and gene names, and chose STAGATE for spatial-domain analysis and BSP for SVG detection. The selected clustering method could run without a GPU, histology image or raw 10x files; the SVG method accepted H5AD counts without a single-cell reference. The workflow identified seven domains across 4,910 spots and 2,325 significant SVGs among 18,785 tested genes, returning domain maps, UMAP embeddings, SVG spatial maps, marker heatmaps and domain-level summaries.

#### 4. Workflow assembly for user-provided 3D MERFISH data

The 3D MERFISH[89] case tests workflow assembly on complex external data (Fig. 4). Three-dimensional spatial transcriptomics is not a larger single-slice problem. Cells are distributed across sections, and their x-y-z coordinates must remain consistent through conversion, filtering, alignment and visualization, with methods that model tissue organization in 3D rather than treating coordinates as extra features. The user supplied a CSV file and requested analysis of single-cell-resolution, multi-slice MERFISH measurements while leaving the method choice open. As a general-purpose baseline, Biomni A1[119] built an AnnData object, stored 2D and 3D coordinates, reduced a 73,655-cell by 156-gene matrix to 50 principal components, formed a 53-dimensional joint embedding, swept five K-means and five Leiden[120] solutions, and selected one by silhouette score (silhouette 0.191), a generic 3D-aware clustering workflow. ST-Coscientist instead treated the request as multi-slice workflow assembly. It read and schema-detected the CSV, scanned animals, wrote standardized .h5ad files, filtered the subset, preserved the raw 3D coordinates, verified their structure, performed multi-slice alignment-aware preparation, and assembled four spatial-domain methods, namely CellCharter[3], PRECAST[19], SPACEL[45] and STAGATE-3D[77]. Each was run with 3D plots, differential-expression summaries and domain-composition heatmaps, and compared against the published annotation by adjusted Rand index and normalized mutual information. CellCharter gave the strongest agreement (cell-class ARI 0.278, NMI 0.344) and was carried forward for marker-based human interpretation. Published annotations guided selection of the result for expert review; the reported agreement scores therefore characterize this exploratory analysis rather than an independent evaluation of the selected method.
